# PRINCIPLES GOVERNING ENDOTHELIAL CAVEOLAE ORGANIZATION DURING ANGIOGENESIS

**DOI:** 10.64898/2026.03.27.714916

**Authors:** Drew B. Grespin, Jasper S. Farrington, Talen G. Niven, Liam J. Russell, Dinah Loerke, Aaryn J. David, Margaret S. Grespin, Chris M. Culkin, Adella P. Bartoletti, Stryder M. Meadows, Erich J. Kushner

**Author notes:** Co-authors (Drew B. Grespin, Jasper S. Farrington). Author for correspondence: Erich J. Kushner, University of Denver, Department of Biological Sciences, Denver, CO 80210.

## Abstract

Caveolae, flask-shaped membrane invaginations highly enriched in endothelial cells, play a central role in buffering membrane tension, yet the principles governing their spatial organization remain elusive. This investigation sought to generate the most comprehensive and systematic analysis of blood vessel caveolar spatial organization. To do so, our group leveraged micropatterning technologies to impose precise biophysical constraints on endothelial cell geometry to probe how caveolae are organized under defined tensional and polarity environments. These experiments were integrated with a high-throughput spatial cell mapping computational pipeline for analyzing thousands of caveolae, providing an extremely high-fidelity analysis. Our results provide a governing framework of how total cellular caveolae are spatially organized during random and directional migration, non-motile polarized, nascent and stable monolayers with differing confinement levels as well as in angiogenic vasculature in vivo. Broadly, our results demonstrated caveolae preferentially organized in the rear of migrating and polarized endothelial cells. In differing monolayer configurations, caveolae default to a peri-junctional spatial organization. Lastly, in mouse retinal blood vessels caveolae are most prominent in the vascular front due to their responsiveness to vascular endothelial growth factor signaling. Overall, these results strongly suggest that caveolae cellular arrangement and number are highly predictive of vascular stability and remodeling states.

## INTRODUCTION

Early in development, endothelial cells sprout, proliferate and migrate to form the primitive blood vessel structures that are later fortified with other cell types. As such, endothelial cells play a dual role in initiating early blood vessel development and postnatally lining the interior of all blood vessels. This requires endothelial cells to demonstrate remarkable morphogenic plasticity, switching between creating thousands of kilometers of vasculature during angiogenesis then converting to an ultra-resilient, low turnover, liquid barrier with extreme longevity. Furthermore, it suggests that many endothelial components may also serve a dual role in developmental and maturation processes. One such component is caveolae, which are flask-shaped plasma membrane invaginations found in nearly all tissue types. Endothelial cells are characterized by the highest enrichment of caveolar structures compared with all other tissue depots^1, 2^. While the genetic ablation of caveolae formation does not affect early survival^3, 4^, loss of caveolae was found to specifically promote negative cardiovascular outcomes leading to early cardiac stiffening and increased incidence of dilated cardiomyopathy^5–7^. A paradox exists mechanistically, as caveolae via Caveolin-1 (Cav1) inhibit endothelial nitric oxide synthase (eNOS), the enzyme responsible for production of nitric oxide (NO) and its myriad of vasodilatory and anti-inflammatory effects^8^. As a result, reduced caveolae numbers augment endothelial NO production and its well-characterized atheroprotective signaling, suggesting a benefit from having fewer caveolae in blood vessels. Additionally, newer evidence suggests the cardiovascular benefits from reduced caveolae could also stem from its role in low-density lipoprotein transport, independent of its interactions with NO production^9^. In development, the ablation of caveolae reduced angiogenic sprouting behaviors^8^. These data suggest that although caveolae are not required for blood vessel formation, they are necessary for normal physiological development.

At nanoscale, caveolae are 70nm pits which are formed through enhancing membrane curvature facilitated by integral membrane proteins, such as Caveolin-1,2,3 and Cavins, that aggregate in cholesterol rich lipid rafts^10^. Much of the early investigations into caveolar function centered on the idea that caveolar pits assist endocytosis. However, new evidence suggests caveolar pits play an important role in mechanotransduction^11, 12^. Notably, several investigations demonstrated that caveolae provided both plasma membrane tension sensing and buffering system, particularly in tissues that are exposed to high levels of mechanical strain^13, 14^. For example, strain exerted on the plasma membrane causes flattening of the caveolar pits back into a planar configuration providing extra membrane, buffering the strain^14^. As caveolae can provide a means of mechanoprotection they tend to reside or cluster next to cell-cell contacts in close proximity to adherens junctions as observed in epithelial sheets^15^. In migrating cells, caveolae form in the rear of the cell^16^. This spatial arrangement of caveolae clustering is deferential to the plasma membrane tensional environment. In other words, caveolae pits form in areas of low membrane stretch, such as on the retracting end of migrating cells^11^. This low-tension biogenesis model is somewhat incomplete as caveolar proteins have been purported to be linked to cytoskeletal structures such as filamentous actin through proteins like EHBP1^17–19^. Therefore, caveolar pits are not completely unencumbered to flow throughout the plasma membrane as they can be tethered to non-membranous structural elements. This calls into question how caveolae are organized in various biological situations. Is caveolar spatial organization ancillary to cell shape and downstream membrane tension or do underlying cytoskeletal elements influence their localization? It can be argued that these are important questions as endothelial caveolae number and location are strongly correlated with atherosclerotic disease progression^9^.

Beyond caveolae showing elevated numbers in atherosclerotic lesions there is not a comprehensive characterization of how caveolae are spatially organized in various single-cell and multi-cell environments in endothelial tissue. Our goal was to describe how caveolae are organized in endothelial cells in various cellular and geometric configurations. Using novel micropatterns, conventional cell culture and in vivo imaging, we identify how cellular arrangements influence caveolar organization in endothelial cells. Leveraging the reproducible shapes imparted by our micropatterning platform, caveolae clusters were analyzed using spatial cell mapping^20^ for high-accuracy representation of puncta locality. Our results showed that individual migrating endothelial cells demonstrated a canonical caveolar morphology in which caveolae cluster in the rear on the trailing membrane. Intriguingly, cellular confinement enhanced unipolar anisotropy which further augmented caveolar rearward localization. Fabricating planar cell polarity-blunting geometries further underscored this finding. In monolayers, caveolae demonstrated temporal recruitment to peri-junctional locations. However, distorting cellular arrangements from interdigitated to head-to-head orientations changed how caveolae localized, likely due to the altered plasma membrane tensional environment. Lastly, we explored how caveolae are arranged during retinal angiogenesis. Our data indicates that caveolae are most highly expressed in the vascular front and their expression precipitously drops in mature vessel beds. Additionally, we demonstrated that Cav1 expression is coupled to VEGF signaling. Overall, our data provides a first of its kind high-definition characterization of caveolar spatial distribution and regulation in blood vessels.

## RESULTS

Our first goal was to explore how caveolae are organized in single, freely migrating endothelial cells. Localization of caveolae structures were marked using a caveolin-1 (Cav1) antibody that was previously shown to be a reliable marker of caveolae location^21^. Endothelial cells (ECs) were also probed for F-actin to delineate the cell perimeter, lamellipodia and general actin morphology. Staining caveolae in randomly migrating ECs did not reveal a discernable localization pattern. Broadly, Cav1 puncta were located towards the interior cell parenchyma being devoid near the lamellipodial projections (**Fig. S1A**). We measured the distance to the cell exterior vs Cav1 intensity (**Fig. S1B**) to more thoroughly scrutinize for localization patterns. Our analysis showed a slight bias of Cav1 puncta toward the interior of the cell (**Fig. S1C**). Presupposing that the lack of planar cell polarity was causing the absence of a defined cellular locality of caveolar clustering, we created a random cell polarity culture environment. Using a circle micropattern geometry, we prevented the cell from elongating in a defined, unidirectional axis; thus, blocking establishment of a front-rear cell polarity (**Fig. 1A**). Circle micropatterned cells once again demonstrated Cav1 localization primarily towards the interior of the cell with a marked exclusion zone near the cell’s outer perimeter (**Fig. 1B**). Given that the Cav1 antibody does not distinguish between endosomal or plasma membrane bound species, we were concerned that some of the perinuclear signal was simply due to Cav1 accumulation in the Golgi apparatus. To address this, we co-stained for Golgi complex marker GalT and Cav1 to assess overlap in signals. Our results suggest very little of the Cav1 staining is associated with its transit through the Golgi complex (**Fig. S1D,E**). To glean a more quantitative representation of Cav1 distribution, we employed spatial cell mapping (SCM)^22^. This approach is largely made possible by leveraging the stereotyped cell shape imparted by the micropattern where numerous cells can be aligned to make an ‘average’ cell^23^. This average cell represents the mean spatial organization of Cav1 puncta localization across multiple cells. Once constructed, probabilistic maps accurately illustrate population-based spatial organization of caveolar puncta. Caveolae showed a clear spatial preference accumulating centrally in the cell (**Fig. 1C,D**). Cav1 plots demonstrated a linear positive relationship where caveolae density is low at the cell perimeter and increases near the cell center (**Fig. 1C,E**). As a non-caveolae comparison, we mapped F-actin localization. Unlike caveolae, F-actin demonstrated a bimodal distribution in which it was most accumulated at the cell cortex and to a lesser extent at the cell center (**Fig. 1C,E**). Overall, these results demonstrate that in a non-polar EC, caveolae maintains a centralized spatial preference.

**Figure 1.**
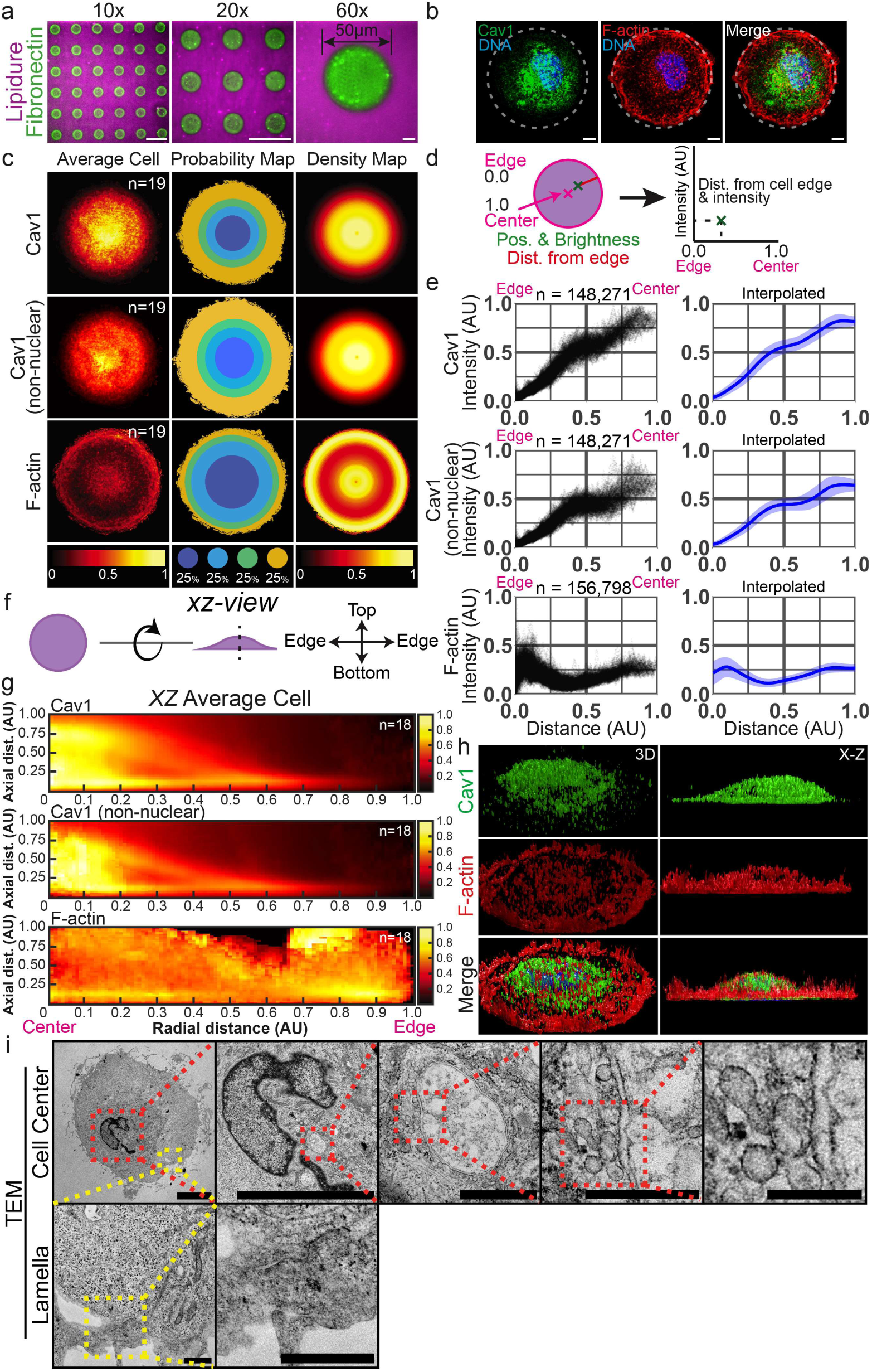
Non-polar endothelial cells show centralized caveolar organization. **(A)** Representative images of circle micropatterns with cell-repulsive Lipidure (magenta) and extracellular matrix (ECM) protein fibronectin (green). Scale bars indicate 100µm (10x & 20x) and 10µm for the 60x image. **(B)** Representative images of endothelial cells stained for Caveolin-1 (Cav1, green), F-actin (red), and DNA (blue) on circle patterns. **(C)** Average cell projection (left), quartile probability map (middle), and radial density map (right) of indicated groups. Intensity and color scales are displayed at the bottom. N= number of cells. **(D)** Cartoon to view the process of distance vs. intensity plotting. **(E)** Distance vs. intensity graph for indicated groups. Distances are calculated and normalized from the edge (0) to the center (1); intensities are also normalized. N=number of points. Interpolated graphs (right panels) show mean line (dark blue) and standard deviation (light blue). **(F)** Cartoon to view the process of distance vs. intensity plotting. **(G)** X-Z view average models for indicated groups. N= number of cells. **(H)** 3-dimensional renderings of Cav1 (top, green), F-actin (middle, red), and a merge including labeled DNA (bottom, blue) from both a 3D (left) and X-Z (right) point of view. **(I)** Transmission Electron Microscopy (TEM) representative images depicting in regions of interest. In the Caveolae row, scale bars indicate 10 µm (left 2) 1µm (middle, 3), 500 nm (4), and 200 nm (rightmost, 5). In the Lamella row, scale bars indicate 1 µm.

Using a similar computational framework, we next aligned circle micropatterned cells in the axial orientation to provide an ‘average cell’ profile to visualize caveolar spatial patterning (**Fig. 1F**). Like the X-Y orientation, Cav1-positive puncta were most intense near the cell center and significantly diminished towards the cell periphery (**Fig. 1G**). Notably, caveolae density was also enriched in the X-Z profile which was confirmed by rendering 3D projections (**Fig. 1H**). This suggests a potential biophysical mechanism that may be enforcing a more centralized caveolar spatial patterning in non-polar, non-motile cells. Scrutinizing the circle micropattern data, F-actin localization is fundamentally exclusionary for caveolar clustering. This observation is congruent with the idea of areas of high-protrusive activity promoting plasma membrane strain and dissolution of caveolar pits. To determine if the nucleus impacted caveolar organization, we digitally removed it from analysis. Exclusion of the nucleus led to a tighter clustering of caveolar signal, confirming a centralized localization (**Fig. 1G**). To further validate our observations, we performed transmission electron microscopy (TEM) analysis in circle micropatterned cells. In line with our Cav1 staining and quantification, we found a paucity of caveolae in the cell’s perimeter (**Fig. 1I**). Our computational analysis suggested an accumulation of Cav1 towards the cell parenchyma. We found plentiful caveolae-like structures which were the correct size range (50-100nm^1^) for internalized caveolae as well as demonstrated more complex fusion morphologies such as rosettes^24^ (**Fig. 1I**). Overall, these results indicate that in a non-polarized cell state caveolae are principally devoid from the cell perimeter and default towards the cell center.

We next induced a strong unidirectional cell polarity by constraining endothelial cells on a line-shaped micropattern (**Fig. 2A**). We fabricated line micropatterns with varying widths (5-20µm) to provide a spectrum of cell constraints; a 5µm width being highly constrained while a 20µm width inducing low-constraint (e.g. enhanced cell spreading) (**Fig. 2B**). The idea behind this approach is that a highly constrained geometry would produce the more extreme cell anisotropy and the greatest commitment to a singular direction. Line micropatterned ECs demonstrated an elongated morphology and a single-sided centrosome orientation in the direction of cell migration (**Fig. 2C**; **S2A,B)**. These results suggest that all micropattern line widths elicited a strong unidirectional polarity.

**Figure 2.**
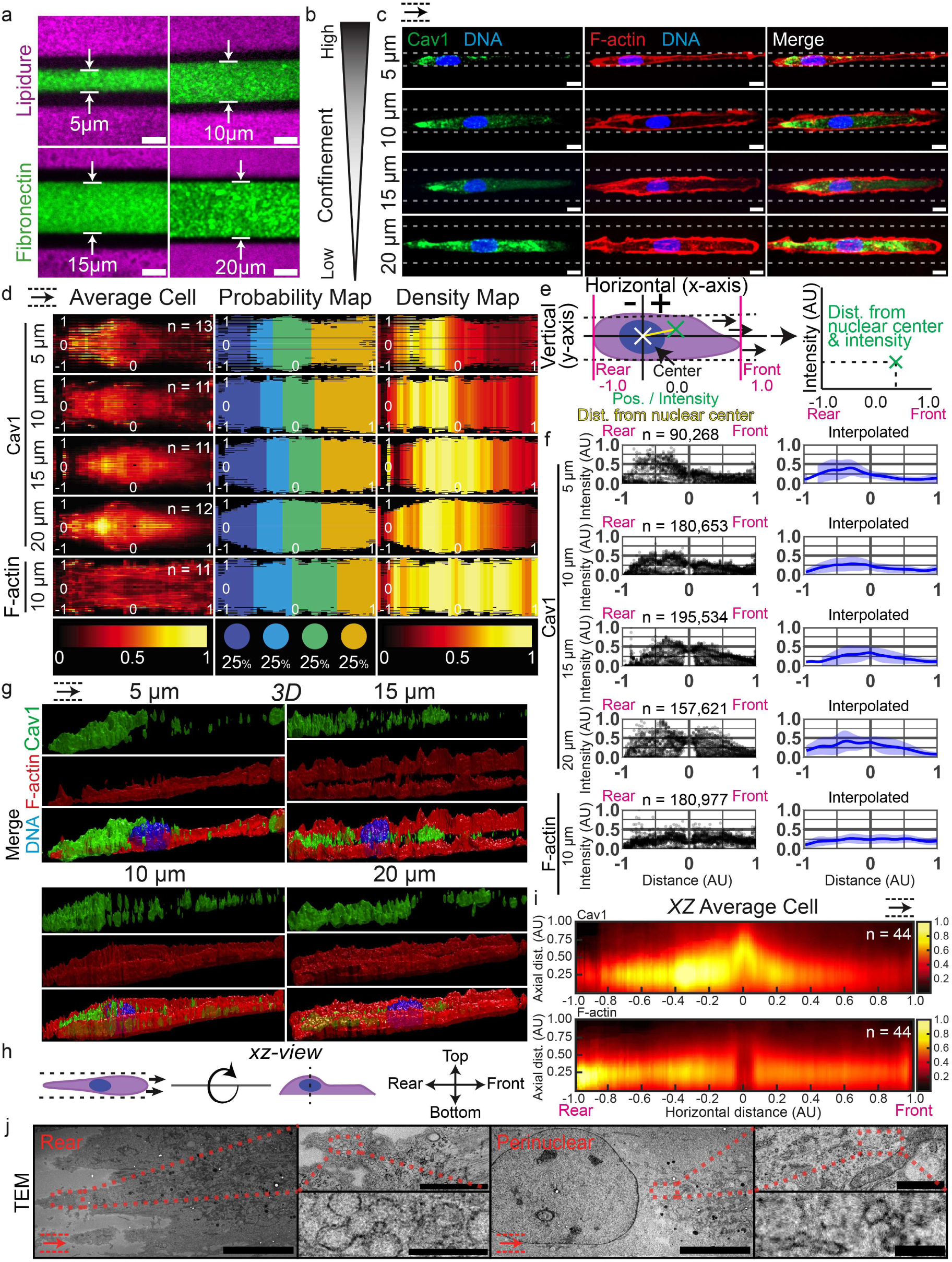
Unidirectional confinement promotes caveolae localization to the retracting end of migrating endothelial cells. **(A)** Line micropatterns of varying widths with cell-repulsive Lipidure (magenta) and extracellular matrix (ECM) protein fibronectin (green). Scale bars represent 10 µm. **(B)** Gradient displaying the transition from low confinement (bottom) to high confinement (top) as line micropattern width decreases in panel C. **(C)** Representative images of migrating endothelial cells stained for Caveolin-1 (Cav1, green), F-actin (red), and DNA (blue) along line patterns. Black arrow indicates direction of migration. **(D)** Average cell projections (left), quartile probability maps (middle), and radial density maps (right) of Cav1 or actin (control) on line patterns with indicated confinement widths. Intensity and color scales are displayed at the bottom. Maps display rightward migration indicated by arrow at the top-left corner. N= number of cells. **(E)** Quantification schematic illustrating distance vs. intensity plotting. **(F)** Distance vs. intensity plots of Cav1 on indicated line patterns. Distances are calculated and normalized from the center (0) to the frontmost (+1, on right) or rearmost points (-1, left); intensities are also normalized. N=number of points. Interpolated graphs (right panels) show mean line (dark blue) and standard deviation (light blue) **(G)** 3-dimensional renderings of Cav1 (top, green) F-actin (middle, red), and a merge on differing width line micropatterns. **(H)** Cartoon displaying the rotation into the X-Z axial view for the average models of Cav1 (top) and F-actin (bottom) shown below. Color scales are shown to the right along with x distance (bottom) and z distance (left). **(I)** X-Z view average models for indicated groups. N= number of cells. **(J)** Transmission Electron Microscopy (TEM) representative images depicting rear and perinuclear region. In the rear (left three) images, scale bars indicate 10µm (large, leftmost) 1µm (small, top), 200nm (small, bottom). In the frontward perinuclear (right 3) images, scale bars indicate 10µm (large, leftmost), 1µm (small, top), and 500nm (small bottom).

Although cells were geometrically constrained via the micropattern, the ECs were only bounded in the Y-dimension; inherently, this led to more shape variability in the longitudinal X-axis as compared to cells constrained in all directions (e.g. circle micropatterns). To account for the increased variability in cell shape in our analysis, we normalized the horizontal distances to the nuclear centroid as a common fiducial marker and plotted caveolae distances and intensity relative to that position (**Fig. 2E**). Our analysis indicated a strong rearward preference of caveolae similar to previous reports^25, 26^(**Fig. 2D**). Again, using F-actin as a non-caveolar comparison, we observed a more homogenous protein distribution (**Fig. 2D**). Similarly, average cell renderings in the axial profile demonstrated a defined rearward spatial patterning preference of caveolae as compared with F-actin (**Fig. 2h,I**). Investigating the dynamic nature of caveolae localization in this environment, we next live-imaged ECs during line migration. Again, live cell dynamics strongly echoed our density mapping fixed cell analysis in which caveolar localization was rearward (**Fig. S2D,E**).

Plotting relative caveolae locations between the different levels of confinement showed a trend in which 5µm lines demonstrated the greatest rearward clustering, which was incrementally reduced with decreasing confinement conditions to 15µm (**Fig. 2F; S2C**). Interestingly, the 20µm, lowest confinement condition, showed high rearward Cav1 clustering (**Fig. 2F**). Perhaps, this is due to more efficient migration in this confinement condition. To better visualize caveolar distribution between confinement groups, we rendered 3D images of caveolae and F-actin (**Fig. 2G**). To confirm this observation, we sectioned through the retracting membrane and found plentiful caveolae structures (**Fig. 2J**). Overall, this data provides strong evidence that caveolae reside in the rearward portion of a unidirectionally polarized, motile EC. Furthermore, there is a strong positive association between the strength of caveolar polarization and cell confinement.

During cell migration the rear membrane physically retracts as focal adhesions connecting the cell to the underlying stratum release. Thereafter, the plasma membrane rebounds, due to release of viscoelastic tension, acutely reducing local membrane stretch. The most current model of caveolae formation suggests that caveolar invaginations preferentially assemble in areas of low membrane tension. However, this observation begs the question, how would caveolae organize in a cell that is not motile, but polarized? Does cell retraction unambiguously dictate caveolae localization, or are there interactions with cell polarity systems? In this scenario, there would be a clear forward-rear cell polarity, but plasma membrane retraction would be absent, limiting local release of membrane tension in the rear of the cell. To test this, we fabricated a crossbow micropattern that enforces a strong planar cell polarity axis in the absence of migration (**Fig. 3A**)^27^. To assess planar cell polarity, we quantified the nuclear-centrosome angle as a proxy for cell orientation as we have previously published^28^. Our average nuclear-centrosome angle of ∼17° (0 ° being directly forward) demonstrates the bulk of nuclear-centrosome angles were directed towards the outer bow region of the micropattern (**Fig. S3A**). This data suggests crossbow micropattern ECs are polarized towards the outer portion of the crossbow micropattern and therefore have a defined polarization axis in the absence of migration. In directionally polarized ECs, we again stained for caveolae and F-actin for SCM. Our results showed a generally broader distribution of caveolae patterning as compared with line micropatterns or the F-actin control; however, there was still a slight, but significant, rearward spatial preference (**Fig. 3B-D**). Using the probability map, approximately 70% of the caveolae is in the bottom half of the cell. In the axial view, Cav1 staining is marginally more prominently in the rearward half of the EC as compared with the front (**Fig. 3F,H**). F-actin control comparison demonstrated a homogenous localization on the leading and lagging ends (**Fig. 3E,H,I**). Intensity plots and forward/rear ratio comparisons for caveolae distribution echoed this observation in which there was a modest rearward localization (**Fig. S3B,C)**. Akin to circle patterns, a prominent Cav1 exclusion zone near the forward-facing bow was also observed, the likely location of lamellipodia and associated branched actin assembly (**Fig. 3C-E**). Zeroing in on the bulk of the caveolar signal, we excluded the outer lamellipodial projections (**Fig. 3G**). The resulting axial view more clearly demonstrated a prominent rearward Cav1 clustering as compared to the cell front (**Fig. 3I**). These data suggest that non-migratory, polarized ECs also reinforce caveolae organizing towards the rear of the cell.

**Figure 3.**
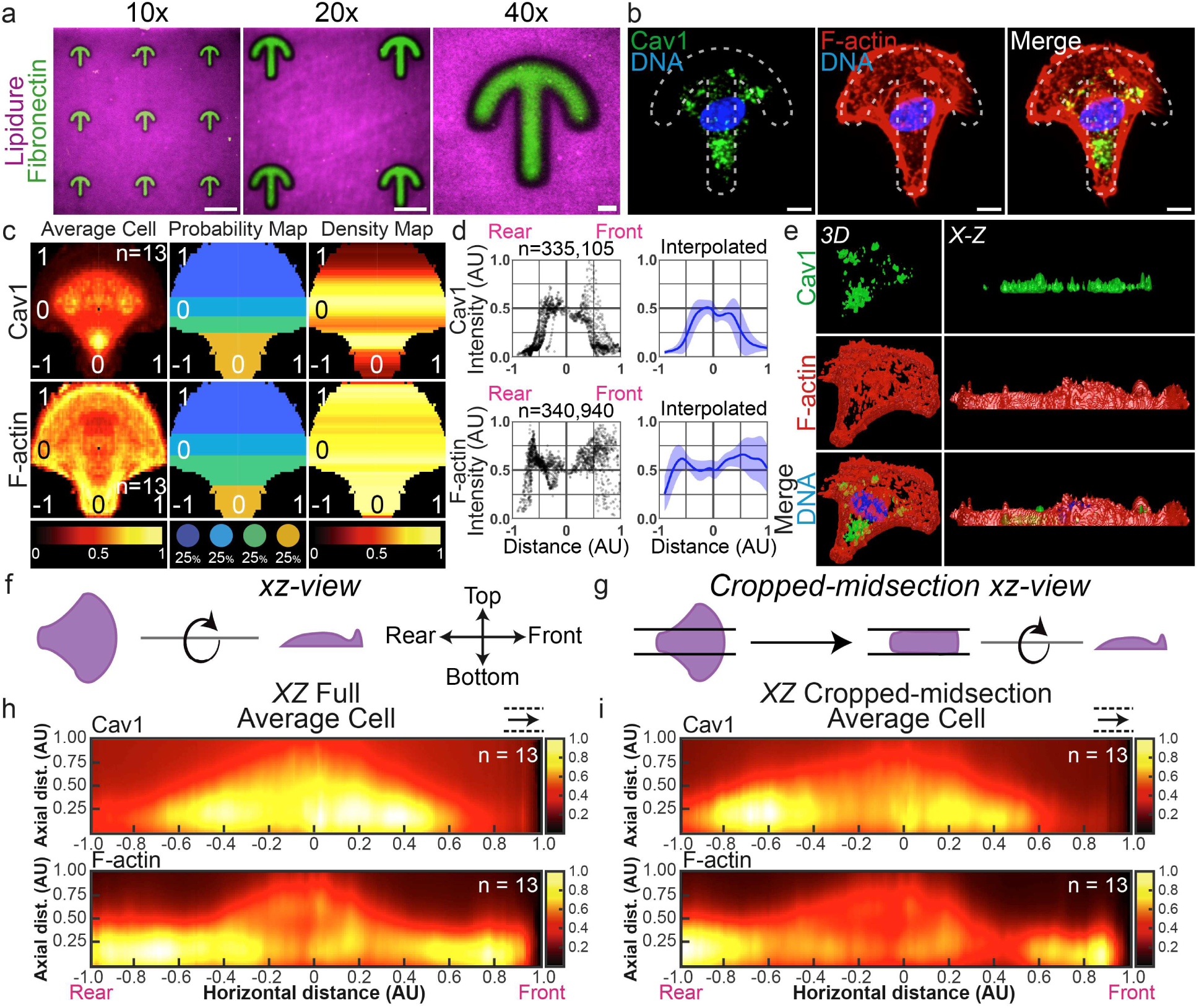
Non-motile, planar-polarized endothelial cells display rearward caveolar localization. **(A)** Images of crossbow micropatterns with cell-repulsive Lipidure (magenta) and extracellular matrix (ECM) protein fibronectin (green). Scale bars represent 10µm. **(B)** Representative endothelial cells stained for Caveolin-1 (Cav1, green), F-actin (red), and DNA (blue), and imaged on crossbow patterns. **(C)** Average cell projections (left), quartile probability maps (middle), and radial density maps (right) of Cav1 (top) and F-actin (bottom) on crossbow patterns. Intensity and color scales are displayed at the bottom. N= number of cells. **(D)** Distance vs. intensity graphs of Cav1 (left) and F-actin (right). Distances are calculated and normalized from the center (0) to the frontmost (+1, right) or rearmost points (-1, left); intensities are also normalized. N=number of points. Interpolated graphs (right panels) show mean line (dark blue) and standard deviation (light blue). **(E)** 3-dimensional renderings of Cav1 (top, green), F-actin (middle, red), and a merge including labeled DNA (bottom, blue) from both a 3D (left) and X-Z (right) point of view. **(F)** Cartoon displaying the rotation into the axial X-Z view for the below X-Z view average models for Cav1 (top) and F-actin (bottom). **(G)** Cartoon displaying cropping the midsection of the average cell and then the same rotation into the X-Z view as F. **(H)** X-Z view full cell average models for indicated groups. N= number of cells. **(I)** X-Z view cropped-midsection cell average models for indicated groups. N= number of cells.

As ECs do not exist as single-celled units in physiological development, we next surveyed how caveolae are spatially organized in multicellular environments. Employing a classic endothelial monolayer, we stained for Cav1, F-actin, and adherens junction marker VE-cadherin (**Fig. S4A**). Although, caveolae puncta were distributed throughout the cell, there was a slight enrichment at the cell-cell junctions. To better quantify this association, we fabricated ‘track’ micropatterns **(Fig. 4A).** The goal was to create a geometry that could be homogenously saturated to maximal confluency. Additionally, the circle configuration inhibited ECs from extraneous migration, allowing ECs to more quickly ‘interlock’ into a stable monolayer confirmation. Lastly, the 50µm width of the track adhesion surface mimicked the diameter of a small capillary, reinforcing a more physiological EC density (**Fig. 4B**). To explore caveolar junctional localization, we performed a time course analysis ranging from 4-24 hours. We observed that Cav1 was initially more dispersed; however, by 8 and 24 hours caveolae puncta were more proximal to cell-cell junctions (**Fig. 4B**). Density plots demonstrated a generally high-level of variability between conditions; however, a modest trend emerged at 24 hours demonstrating increased peri-junction caveolar density (**Fig. 4C,D**). To explore this junction accumulation further, we binned caveolar puncta into junctional (proximal to cell perimeter) and the remaining pool (non-junctional) between incubation times (**Fig. 4E**). A trend of more junctional Cav1 puncta at 24hrs as compared to the lower incubation times was observed (**Fig. 4F, S4C-D**). Lastly, we confirmed the presence of caveolae pits at the cell-cell interface with TEM imaging (**Fig. 4G**). These data indicate in confluent monolayer, caveolae position in a peri-junctional configuration.

**Figure 4.**
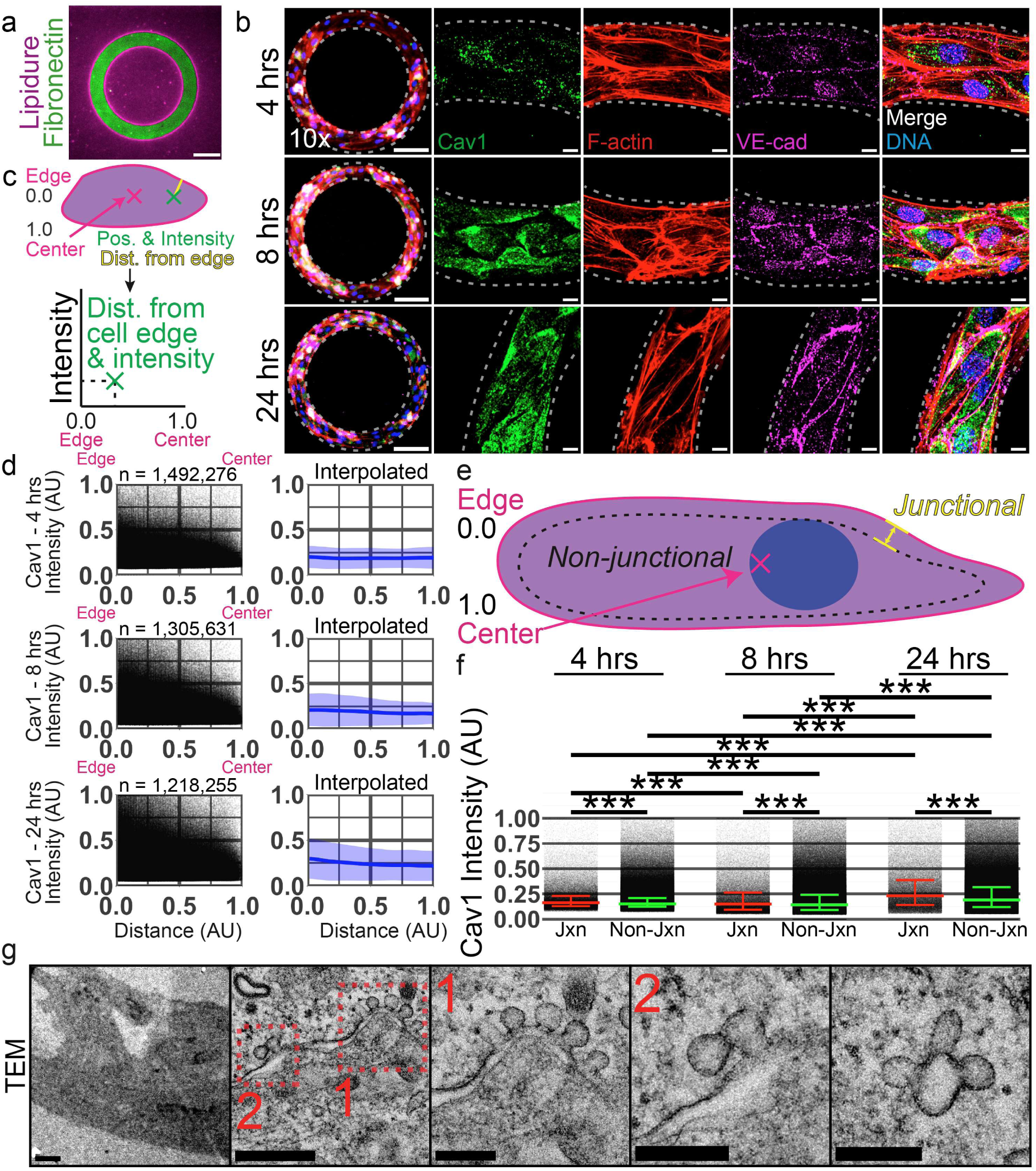
Junctional stability is associated with caveolae in close proximity to cell-cell contacts. **(A)** Track micropatterns with cell-repulsive Lipidure (magenta) and extracellular matrix (ECM) protein fibronectin (green). Scale bar represents 100µm. **(B)** Cartoon displaying the distance vs. intensity graphs shown in panel D. **(C)** Representative endothelial cells imaged on track micropatterns incubated for 4 (top), 8 (middle), and 24 (bottom) hours (hrs) and stained for vascular endothelial cadherin (VE-cad, magenta), F-actin (Actin, red), Caveolin-1 (Cav1, green). Insets are areas of higher magnification. Scale bars represent 50 µm for the 10x images and 10µm for the 60x images. **(D)** Distance vs. intensity graphs showing quantification of distance from the junction (0) to the center of the cell (1) on the horizontal axis and normalized Cav1 intensity on the vertical axis for indicated times. N=number of points. Interpolated graphs (right panels) show mean line (dark blue) and standard deviation (light blue). **(E)** Schematic of junctional and non-junctional zones quantified in panel F. **(F)** Non-junctional vs. Junctional bar chart of Cav1 intensity at the indicated time points. Number of points per condition is as follows: 4 hour Junctional = 110,813; 4 hour Non-Junctional = 1,381,463; 8 hour Junctional = 91,082; 8 hour Non-Junctional = 1,214,549; 24 hour Junctional = 84,474; 24 hour Non-Junctional = 1,133,781. **(G)** Transmission Electron Microscopy (TEM) representative images depicting junctional regions on track micropatterns. In the rear (left three) images, scale bars indicate 10µm (large, leftmost) 1µm (small, top), 200nm (small, bottom). In the frontward perinuclear (right 3) images, scale bars indicate 10µm (large, leftmost), 1µm (small, top), and 500nm (small bottom). ***P<0.001. Additional statistical comparisons in supplemental data.

We next explored how confinement influenced caveolar spatial patterning in multicellular scenarios. Again, we patterned EC monolayers on various line widths. Interestingly, at high confinement (e.g. 5µm width) ECs were arranged in a head-to-tail orientation, similar to those morphologies described in small connecting capillaries^29^(**Fig. 5A)**. Decreased confinement promoted the interdigitation of ECs with much of the cell perimeter contacting its neighbors (**Fig. 5A**). Due to the head-to-tail orientation, we excluded 5µm widths from analysis as there were insufficient junctional contacts to analyze. Binning the remaining line widths (10-20 µm) we observed a modest trend with Cav1 being more present near junctional areas (**Fig. 5B-D**). Similarly, there was a modest positive linear correlation between junctional F-actin and Cav1 intensity (**Fig. S5A**). Binning the Cav1 into junctional and non-junctional pools across all widths there was no biologically meaningful difference, although statistically different, in Cav1 locality (**Fig. 5C, S5B**). Further exploring this relationship, we quantified the junctional/non-junctional Cav1 spatial distribution for individual confinement groups. All confinements produced a modest junctional preference (**Fig. 5F, S5C,D**). Similar to track patterns, we confirmed the presence of caveolae structures at the cell-cell contact sites using TEM imaging (**Fig. 5G**). Overall, this data indicates caveolae have a slight junctional preference in non-migratory, confluent, arrangements and cellular confinement plays little part in the spatial distribution of caveolae.

**Figure 5.**
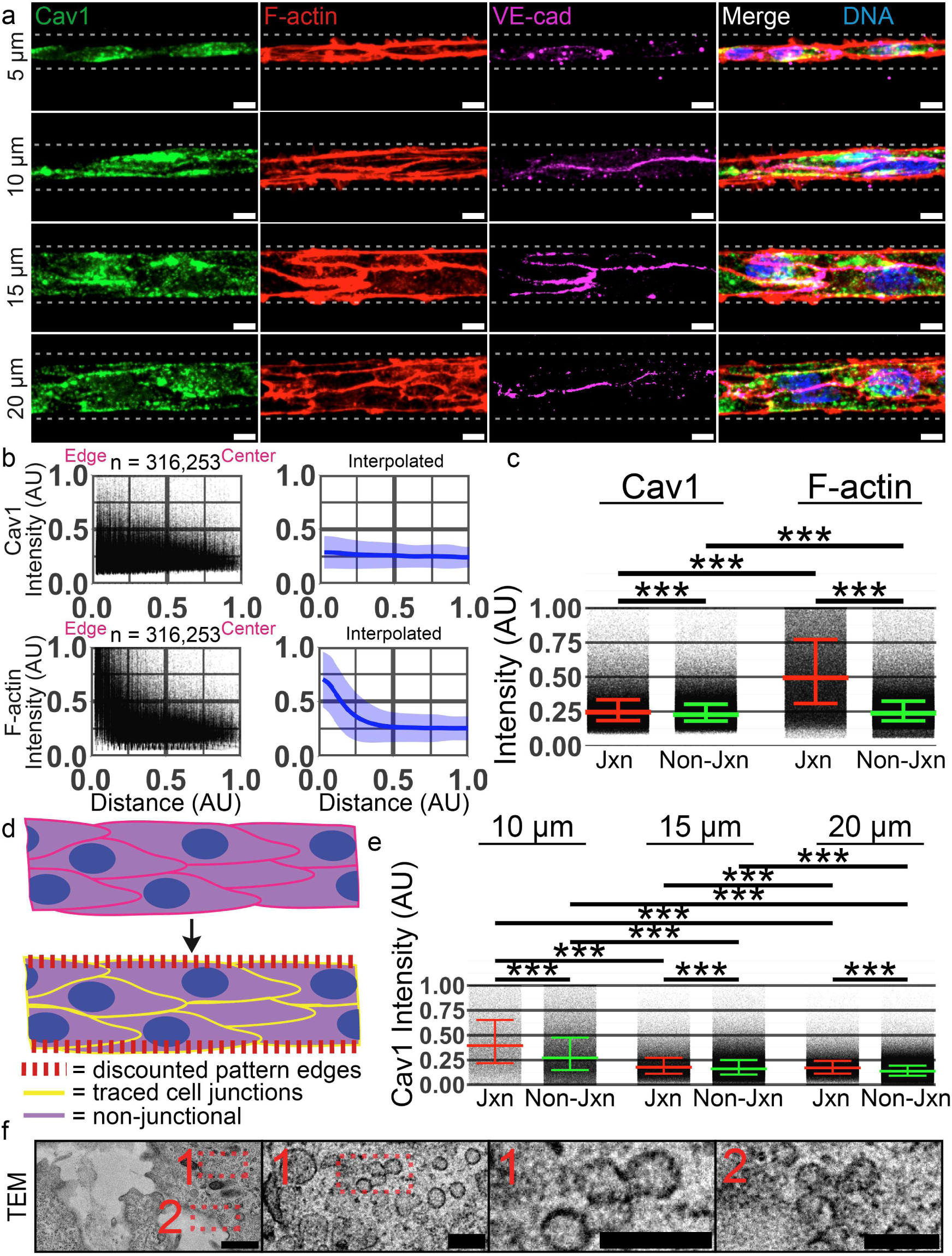
Cell confinement alters caveolar spatial preferences. **(A)** Representative images of confluent, line-patterned endothelial cells fixed 24 hours (hrs) after plating and stained for indicated proteins. Scale bars represent 10 µm. **(B)** 15 µm line distance vs. intensity graphs showing quantification of distance from the junction (0) to the center of the cell (1) on the horizontal axis and normalized Cav1 or Actin intensity on the vertical axis. N=number of points. **(C)** Non-junctional vs. Junctional bar chart of Cav1 intensity for 15 µm line. Number of points per condition is as followed: Cav1 Junctional = 125,904; Cav1 Non-Junctional = 190,349; Actin Junctional = 125,904; Actin Non-Junctional = 190,349. (**D**) Cartoon of junctional quantification approach. (**E**) Non-junctional vs. Junctional bar chart of Cav1 intensity at indicated confinements. Number of points per condition is as follows: 10 µm Junctional = 48,796; 10µm Non-Junctional = 124,317; 15µm Junctional = 120,416; 15µm Non-Junctional = 285,242; 20µm Junctional = 147,039; 20µm; Non-Junctional = 515,550. (F) Transmission Electron Microscopy (TEM) representative images depicting junctional regions on track micropatterns. In the rear (left three) images, scale bars indicate 1µm (large, leftmost) and 200nm for insets.***P<0.001. Additional statistical comparisons in supplemental data.

We next sought to characterize how caveolae are arranged in vivo. To do so, we harvested postnatal (P) day 7 mouse retinas and stained for junctions (VE-cadherin), ERG and caveolae. We were immediately struck by the difference in caveolae expression between the vascular front and distal stable blood vessels. In the migratory front, where angiogenesis is maximally active, there was significantly greater caveolae intensity compared with blood vessels towards the optic nerve (**Fig. 6A**), classically associated with a more stable vascular phenotype^30^. Using SCM, we measured Cav1 intensity and location across the retina (**Fig. 6B**). We observed a clear positive relationship with Cav1 intensity being elevated closer to the vascular front (**Fig. 6C,D; S6A-E**). Assessing the Cav1 distribution in individual ECs, we again observed that Cav1 maintains a rearward spatial preference in angiogenic tips cells (**Fig. 6E,F; S6F,G**). To our knowledge, this is the first report demonstrating a positive association between caveolae expression and areas of angiogenic remodeling. This data suggests that caveolae are most associated with nascent blood vessel growth.

**Figure 6.**
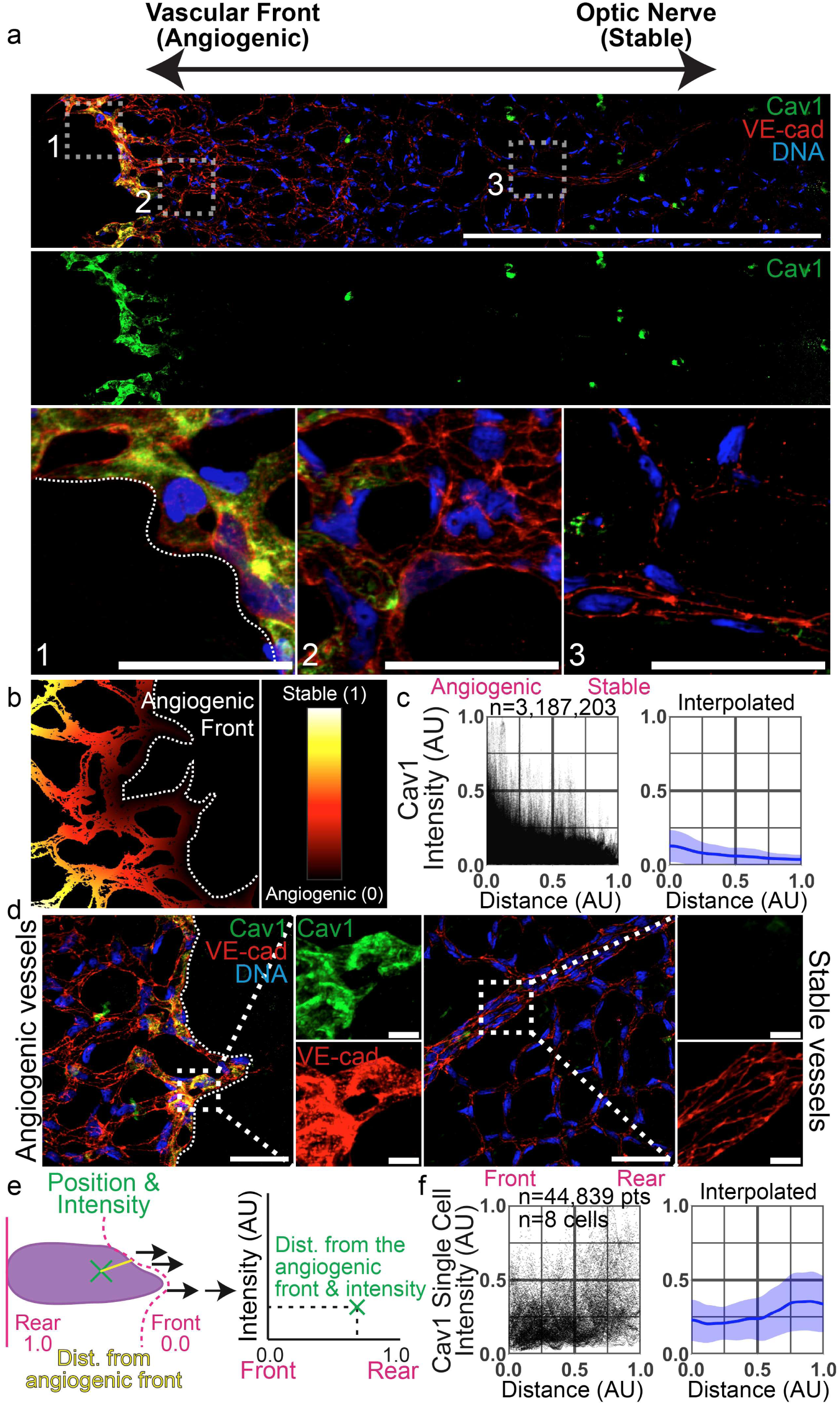
In vivo mouse retinovasculature shows high Caveolin-1 expression at the angiogenic front. (**A**) Post-natal day 7 mouse retina vessel bed stained for Caveolin-1 (Cav1, green, isolated in middle image), endothelial cadherin (VE-cad, red) and endothelial specific nuclei (ERG, blue). Insets 1, 2, and 3 demarcate migratory (1) to most stable (3) are shown at the bottom with scale bars representing 50 µm. (**B**) Image displaying distance of endothelial cells to the angiogenic front (0 being proximal, 1 distal) used for quantification in panel C. (**C**) Distance from the angiogenic front (horizontal axis) vs. normalized Cav1 intensity (vertical axis). N = number of points. n = 3,187,203 pts collected from 10 images with an average of 54 cells per image. Interpolated graphs (right panels) show mean line (dark blue) and standard deviation (light blue). (**D**) Representative images of vasculature at the angiogenic front (leftmost 3 images) and in stable vessels (rightmost 3 images) stained for Cav1 (green, bottom inset), VE-cad (red, top inset), and DNA (blue). Scale bars represent 50 µm and 10 µm in the magnifications shown to the right. (**E**) Quantification schematic illustrating distance vs. intensity plotting for individual endothelial cells. (**F**) Distance from the front of the cell (horizontal axis) vs. Normalized Cav1 intensity (vertical axis) plot. Number of points and cells displayed. Interpolated graphs (right panels) show mean line (dark blue) and standard deviation (light blue).

To explore what triggers caveolae expression, we investigated two potential possibilities. Mechanical tension triggers Cav1 expression. Secondly, Cav1 expression is downstream of growth factor signaling. To test the former, we created a quasi-scratch wound assay in which ECs were plated on a line micropattern with a removable barrier. In a minimal media, containing no exogenous growth factors, we found that ECs migrate into the voided space; however, there was no discernable Cav1 expression, particularly in the leading ECs (**Fig. 7A**). Conversely, performing the same experiment with growth factor supplemented media or with VEGF supplementation there was an increase in Cav1 expression (**Fig. 7A**). Nevertheless, the expression was not confined to leading tip cell or distal stalk cell zones, suggesting a global, unrestricted, increase in Cav1 expression.

**Figure 7.**
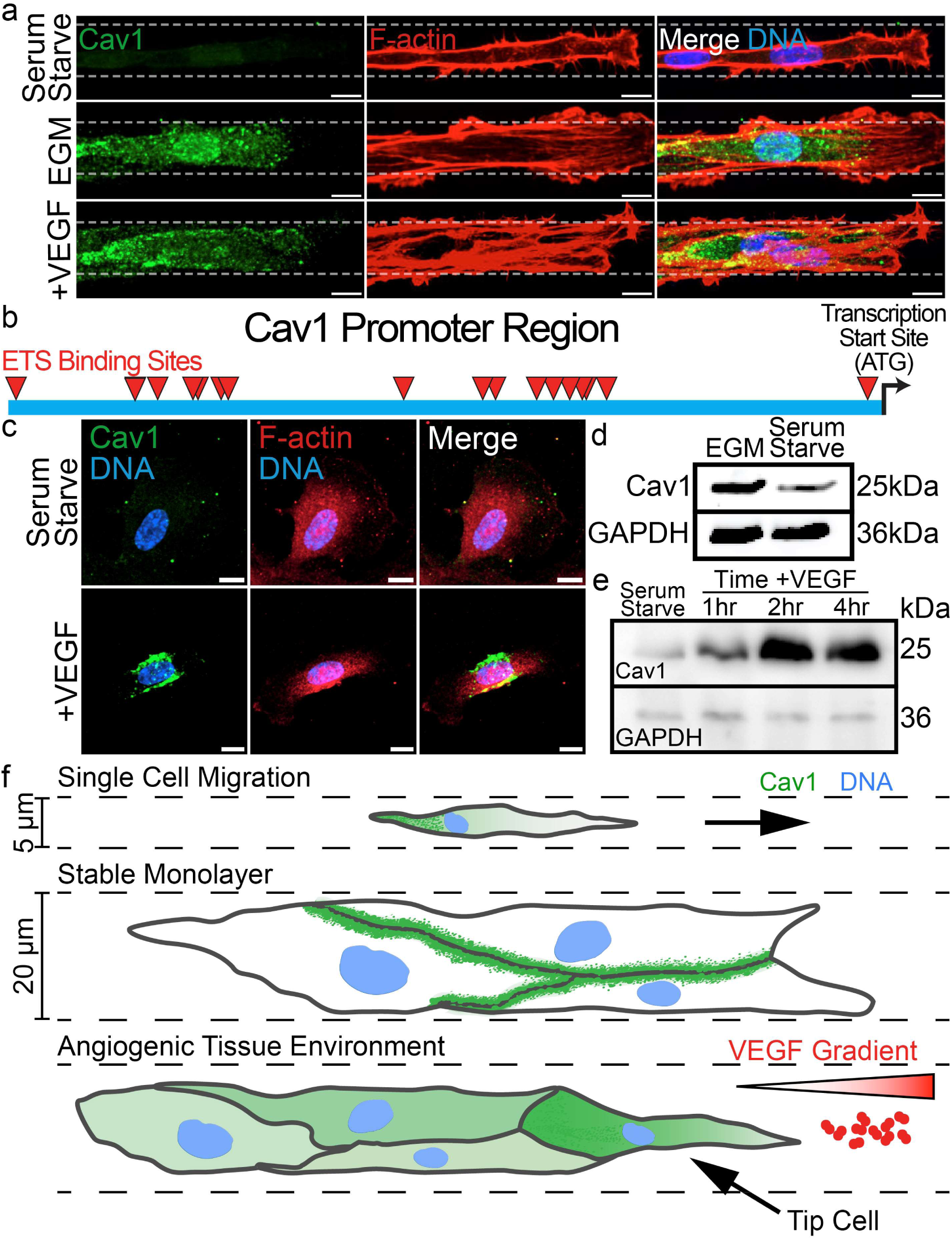
Caveolin-1 protein expression is upregulated by VEGF stimulation. (**A**) Representative images of tip cells (endothelial cells) on scratch wound line micropatterns fixed and stained for Caveolin-1 (Cav1, green), filamentous actin (F-actin, red), and DNA (blue) in different growth factor conditions including EGM (Endothelial Growth Medium), VEGF (Vascular Endothelial Growth Factor) supplementation, and the absence of growth factors (serum starve). (**B**) Schematic displaying the ETS transcription factor binding sites in the upstream promotor region of Cav1. (**C**) Representative images of endothelial cells stained for Cav1 (green), F-actin (red) and DNA (blue) in the presence (top) or absence (bottom, serum starve) of VEGF. Scale bars indicate10 µm. (**D**) Western blot depicting protein expression levels for the same conditions shown in C. (**E**) Western blot depicting protein expression levels of Cav1 (top) and GAPDH (bottom) over the indicated time after VEGF addition. (**F**) Model: Top panel shows endothelial cells seeded in higher confinement (as in the 5 µm line pattern) conditions establish more of a front-rear migratory phenotype, and caveolae are localized to the rear. The center of the model depicts that caveolae are junctionally localized upon junctional stabilization. The bottom panel suggests caveolae are plentiful towards the angiogenic front as VEGF signaling triggers Cav1 expression.

To next test if Cav1 expression was sensitive to classic VEGFR2 signaling, we first analyzed the Cav1 upstream promoter region by scrutinizing for VEGF-signaling responsive elements. We found an abundance of Ets (GGA(A/T)) sequences in the -1500 region (**Fig. 7B**). Ets1/2 is activated by VEGFR2 signaling and are known to modulate gene networks containing Ets sites^31^. Next, we checked for protein expression of Cav1 by either withholding or adding VEGF ligand. Adding VEGF significantly increased Cav1 expression **(Fig. 7C-E)** This data suggests that caveolae are integral components that are upregulated in response to proangiongenic cues.

## DISCUSSION

In this report, we used a combination of micropatterning environments and next-generation image analysis to provide fine grain quantification of how endothelial caveolae behave in simple and complex cellular environments. Our results both confirm and extend previous research demonstrating how caveolae are localized to the trailing end of migrating ECs. Interestingly, this trailing end spatial arrangement is primarily due to motile cells releasing membrane tension in the retraction fibers. Interestingly, in non-motile cells with directional polarity a rearward caveolar localization was also maintained. In multiple micropatterned collective-cell environments, caveolae displayed a positive association with elevated junctional stability suggesting a potential interaction with adhesion proteins or signaling. In a stable monolayer, the majority of Cav1 is juxta-junctional on the plasma membrane participating in caveolae formation. At the tissue level, we were surprised to find that caveolae expression in mouse retinas were almost exclusively restricted to the vascular front (**Fig. 7F**). The mechanism underpinning this observation is that Cav1 and several other caveolae proteins are VEGFR2 responsive, thus biologically tied to angiogenic signaling and blood vessel growth. Overall, this investigation provides both a novel analysis of caveolar spatial organization in endothelial cells and blood vessels as well as new insights into their spatial regulation.

Our original entry point into the current investigation was related to study of a separate caveolar protein EHD2 ^17^. In attempting to assign ‘normal’ and ‘dysmorphic’ caveolar spatial arrangements, it was clear that there was no strong consensus on how caveolae organized in endothelial cells, or other tissues at large. This is not to say, others have not investigated caveolar assembly, our contention is there are no highly quantitative reports specifically detailing native caveolae spatial organization preferences. Generally, caveolae locality demonstrates cell type specificity, in some cases being utterly absent^32^, as well as situational differences. Interestingly, endothelial cells demonstrate one of the highest, if not the greatest, caveolae levels in any tissue system^1^. In many instances protein quantification can be handicapped by punctate localization and generally high cell-to-cell variability. Caveolae and their resident proteins are both punctate and variable in expression and appearance, which may, to some extent, hamper previous efforts into precision mapping. Adding to the complexity, caveolae are greatly influenced by tissue state (e.g. single cell, motile, non-motile, monolayer, etc). The lack of consensus and relation to disease pathology in blood vessel homeostasis prompted our group to devise next-generation particle quantification algorithms to provide a thorough investigation of endothelial caveolae organization.

In terms of mechanical strain, few organs are exposed to more than endothelial cells. The neointimal endothelium experiences persistent oscillatory shear stress and circumferential strain due to the hydrodynamic nature of blood flow. Logically, it would follow that a tissue subjected to elevated strain would compensate by increasing mechanisms that safeguard its integrity, such as caveolar proteins. Our results in mapping caveolar localization in retinal blood vessels were unexpected. Our initial thought was that Cav1 (as a proxy for caveolae themselves) would be more enriched in established, stable vasculature to accommodate the strain associated with blood flow; others have alluded to this type of association^33^. On the contrary, Cav1 was almost exclusively expressed in the remodeling vascular front being largely devoid in the trailing network. The narrow band of Cav1 expression at the vascular front was reminiscent of classic notch signaling restriction of VEGFR2 expression in the trailing stalk cell vasculature^34–36^; although we did not test this directly. Others have reported VEGF signaling triggering Cav1 expression in cell culture^37^. Broadly, this suggests caveolae may not participate in buffering the hydrodynamic forces associated with blood flow, perhaps its primary role is absorbing mechanical strain associated with cell migration and vascular remodeling. The ubiquity of caveolae in most cellular systems is somewhat at odds with its purported dispensability in development. Nonetheless, what is coming into the fore is the necessity of caveolae in proper blood vessel development and homeostasis. Caveolae are more prominent in atherosclerotic and cancerous environments. This observation is in line with our data and others in which the proangiogenic environment (disease or developmental) triggers elevated expression of Cav1 for buffering plasma membrane tension excursions related to vascular remodeling. We contend that mitigating the cellular strain from hemodynamic forces associated with blood flow likely do not require caveolae remodeling; although, we did not directly test this in our study. In this light, reports demonstrating genetic ablation of Cav1 or related proteins are not lethal but do distort angiogenic parameters is congruent with our finding^8, 38^.

Cav1 and caveolae pits are more plentiful in solid cancers^26, 38, 39^, an environment laden with excess growth factors and chronic tissue remodeling. A parallel could be drawn with atherosclerotic lesions, with high local concentrations of inflammatory cytokines and pathologic tissue reorganization. Caveolar proteins general responsiveness to VEGF and other mitogens could account for their increased presence in these disease scenarios^40^. Thus, caveolae number, *per se*, may be used as a proxy of vascular remodeling in developmental and disease states^41^. Equally, the lack of Cav1 expression, or other caveolae proteins, could be symbolic of high-cellular stability.

## ACKNOWLEDGEMENTS

Work was supported by funding from the National Heart Lung Blood Institute (R01HL155921-01A1) (EJK).

## AUTHOR CONTRIBUTIONS

JSF, DBG, TGN, MK, AD, MG and CMC performed all experiments. DBG, LJR, and DL performed computational analysis. AB and SMM performed mouse retina work. EJK, JSF and DBG wrote the manuscript.

## DATA AVAILABLITY

Authors will make any data, analytic methods, and study materials available to other researchers written requests to Dr. Erich Kushner.

## METHODS

All research complied with the University of Denver Institutional Biosafety Committee (IBC).

### Reagents

All reagents, siRNA and plasmid information are listed in the reagents table in the supplementary information (Supplementary Tables 1-5).

### Cell Culture

Pooled Human umbilical vein endothelial cells (HUVECs) were purchased from PromoCell and cultured in proprietary media for 2-5 passages. All cells were maintained in a humidified incubator at 37°C and 5% CO_2_. Small interfering RNA was introduced into primary HUVEC using the Neon® transfection system (ThermoFisher) resuspended to a 20µM stock concentration and used at 0.5 µM. Normal human lung fibroblasts and HEK-A were maintained in Dulbeccos Modified Medium (DMEM) supplemented with 10% fetal bovine serum and antibiotics. For 2-dimensional live-imaging experiments, cells were imaged for one minute at baseline before treatment with CK-666 (1μM), and then imaged for an additional two minutes using 5 second intervals. For ligand-modulated antibody fragments tether to the mitochondria (Mito-LAMA) experiments procedures were carried out as previously described ^42^. All plasmids are listed in the supplementary tables.

### Immunoblotting

HUVEC cultures were incubated at 37 overnight in Serum starved media. The following morning, they were changed to a fresh batch of Serum Starve, Serum Starve with the addition of VEGF (20ng/mL), or Endothelial Growth Media for 6 hours. They were then lysed using Ripa buffer containing Protease Inhibitor Cocktail-and processed.

### Immunofluorescence and Microscopy

For immunofluorescence imaging, HUVECs were fixed with 4% paraformaldehyde (PFA) for 7 minutes. ECs were then washed three times with PBS and permeabilized with 0.5% Triton-X (Sigma) for 10 minutes. After permeabilization, cells were washed three times with PBS. ECs were then blocked with 2% bovine serum albumin (BSA) for 30 minutes. Once blocked, primary antibodies were incubated for approximately 4-24 hours. Thereafter, primary antibodies were removed, and the cells were washed 3 times with PBS. Secondary antibody with 2% BSA were added and incubated for approximately 1-2 hours, washed 3 times with PBS and mounted on a slide for imaging. All primary and secondary antibodies are listed in the Supplemental Data 3. All images were taken on a Nikon Eclipse Ti inverted microscope equipped with a CSU-X1 Yokogawa spinning disk field scanning confocal system and a Hamamatusu EM-CCD digital camera. Images were captured using a Nikon Plan Apo 60x NA 1.40 oil objective using Olympus type F immersion oil NA 1.518, Nikon Apo LWD 20x NA 0.95 or Nikon Apo LWD 40x NA 1.15 water objective. All images were processed using ImageJ (FIJI).

### Micropattern Fabrication

Except for the Lipidure coating, micropatterning fabrication was carried out as previously outlined by Azioune et al.^43^ Briefly, coverslips were plasma cleaned at atmospheric conditions for 20 seconds at 25% power. Afterwards, 50 µL of TI Prime (Microchemicals) was spin-coated onto the activated coverslips and then cured on a hot plate at 120°C for 1 minute. Once cured, 0.5% polystyrene dissolved in toluene was spin-coated onto the coverslips. When creating micropatterned glass coverslips, the TI Prime and polystyrene steps were omitted. To adhere the Lipidure to the polystyrene, the coverslips were plasma cleaned again, and then 75 µL of 0.125% Lipidure dissolved in 100% ethanol was spin-coated onto the coverslips. At this point, coverslips can be stored at room temperature. Prior to deep UV exposure, Lipidure-coated coverslips were briefly soaked in PBS for 10-15 minutes. After rehydration, 35 µL of ddH_2_O was used to adhere the coverslips, Lipidure side down, onto the bottom of a photomask (non-chrome side). The photomask was then placed into a deep UV chamber, chrome side up, for 3 minutes. Following UV exposure, a liberal amount of water was added to the coverslips to dislodge them from the photomask. Lastly, patterned coverslips were placed on a 35 µL fibronectin (2ug/mL in 100mM HEPES, pH8) droplet and incubated for 30min at room temperature to graft fibronectin to the vacant patterned areas. Please see Supplemental Information Table 3 for all reagents associated with fabrication and Supplemental Information Table 4 for a list of required equipment. Also please see the Supplemental Information Extended Procedures for a more detailed step-by-step protocol.

### ADAPT Quantification

To track F-actin cellular dynamics, the ADAPT opensource plugin for FIJI ImageJ created by Barry et al.[12] was employed. The segmented ADAPT distances were imported into GraphPad Prism from the plugin’s output data for graphing.

### Image and data analysis

Image and data analysis were performed in MATLAB. ChatGPT was used to assist in refining analytical techniques and improving workflow efficiency. For all images, the actin channel was used to segment the cell using an adaptive thresholding method developed with the Image Segmenter application. Size thresholding was then applied to remove detected objects outside the biologically relevant range for endothelial cells. Finally, background noise was subtracted from the Cav1 channel to improve accuracy before normalizing each cell to its own maximum value. Background noise was estimated using histogram-based analysis to identify the intensity value at which the pixel count reaches a local minimum, marking the transition between cellular signal and non-cellular background. This was further validated by visually inspecting regions devoid of cells.

After normalization, each experimental condition (micropatterned environment or confluency level) was aligned and analyzed specifically to the condition, which is detailed below, so the data and visualizations could be as consistent as possible across disparate conditions.

### Distance vs. Intensity Plots

Distance vs. Intensity plots were created using a custom MATLAB interpolation and plotting function, but the plots were created in RStudio using the high graphics plotting package “ggplot2”. Interpolated lines were plotted with a ribbon for standard deviation because standard error from the mean was too small to be noticeable.

### Circle pattern alignment and analysis

After normalization circle-patterned cells were aligned to their centroid positions in the actin channel. Then, the mean was taken for each specific (x, y) pixel pair across all cells to create an average cell representation that was used for visualization and analysis.

To generate probability and density maps, the Euclidean distances (*d*) of segmented cell pixels from the cell center were computed and rounded:

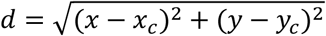

where (*x*, *y*) represents the pixel coordinates and (*x*_c_, *y*_c_) denotes the cell center coordinates. The probability of the protein of interest being located between the center and a given radius—progressively increasing until reaching the cell edge—was determined based on the cumulative sum of its intensity distribution.

To visualize the spatial distribution of the data, an approximated quartile plot was generated, dividing the cell into equally probable regions (each with a 25% probability). The density plots depict regions where the mean intensity is approximately x% of the maximum intensity found in the average cell.

Lastly, Euclidean distances were similarly computed for pixels relative to the cell edge to construct a distance vs. intensity plot described in more detail above.

### Circle pattern 3D alignment and analysis

Using a separate function in MATLAB, cells were thresholded the same as above. Then, the maximum number of Z-slices was calculated across cells. From that number, all cells’ slices were interpolated to be equal to that number, and averages were calculated for each region depth with 0 being the bottom slice and 40 the top. Importantly, these averages were calculated radially, so each radius with respect to x & y away from the center was considered the same for each height. From the center (0, left) to edge (1, right), average normalized intensities across all cells at each normalized radius and height were plotted.

### Single-cell line pattern alignment and analysis

After normalization, line-patterned cells were rotated to be as horizontal as possible. If necessary, they were flipped so that the retracting end was positioned on the left and the leading edge on the right based on actin localization. Cells were then mapped based on their nuclear centroid position.

Within each cell, pixel-wise distances from the nuclear centroid were calculated, including separate horizontal and vertical distances, as well as the combined Euclidean distance. To account for variability in cell length, distances were normalized such that the furthest point in the retracting tail was assigned a value of -1, while the furthest point in the protruding edge was set to +1, but all vertical distances (y) were treated as positive. Finally, distances were binned in both x and y dimensions to generate an average cell representation that could be used for visualization and analysis. Since all vertical distances were treated as positive, mirrored cells were generated in Adobe Illustrator to create a full-cell representation for easier visualization, rather than displaying only half of the average cell. However, the creation of this ‘pseudo-average’ cell was performed post-analysis and did not affect plotting or protein mapping.

Distance vs. intensity plots were generated similarly to those for circular patterns, except that Euclidean distance from the nuclear centroid was used instead of distance from the cell edge to ensure consistency with the visual data from the average cell.

Probability mapping was also performed in a comparable manner, but instead of radiating outward from the cell center, calculations began at the retracting end and progressed toward the protruding end (left to right).

Density visualization followed the same approach as in the circular pattern, where each region’s mean intensity was compared to the maximum intensity value. However, for track-patterned cells, densities were displayed as the average intensity within each *x*-position bin, rather than radial regions.

### Single-cell 3D line pattern alignment and analysis

3D line pattern alignment Z-slice calculation and normalization was performed as above for the circle pattern. The difference in this alignment was that Y-distance from the center line of the nucleus was compressed as non-significant for best averaging. Thus, at one X-distance, every Y-position in one Z-slice was averaged into the same bin. The resulting plot shows the average normalized intensities at each X-position for each interpolated Z-slice across all cells.

### Single-cell crossbow pattern alignment and analysis

The crossbow pattern alignment, segmentation, and analysis closely relates with the principles of the single-cell line cells, except that the crossbow confinement occurs in all dimensions.

### Single-cell 3D crossbow pattern alignment and analysis

The crossbow pattern 3D alignment and analysis also closely relates to the corresponding line pattern analysis except that distances are measured based on the pattern not the nucleus.

### Unpatterned ‘free range’ analysis

Unpatterned cells were segmented using an adaptive thresholding technique similar to that used for the circle-patterned cells. However, an average cell was not generated, as two-dimensional averaging is not feasible for cells with highly variable shapes. However, a distance-from-edge vs. intensity plot was created to analyze these cells like that of the other conditions.

### Track and confluent line pattern analysis

Track-patterned cells were analyzed similarly to unpatterned (’free-range’) cells due to their shape variability. However, cell outlines were manually traced because nuanced junctional boundaries did not allow for automated segmentation.

### Mouse retinovascular analysis

To complete an endothelial-specific analysis of Caveolin-1 staining in mice, we stained C57BL6 P7 mice for Caveolin-1, ERG, a generally known endothelial-specific nuclear marker, and VE-cad. Endothelial cells were identified using ERG as previously reported^44^. For images including tip cells, the angiogenic/migratory front was traced following the tip cells, and then Euclidean distances were calculated from the front (0) to the maximum distance (1) of an endothelial cell in the image. Then the distance vs. intensity was plotted for several images as described above.

### Statistical Analysis

Experiments were repeated a minimum of three times. Statistical analysis and graphing were performed using GraphPad Prism. Statistical significance was assessed with a student’s unpaired t-test for a two-group comparison. Multiple group comparisons were carried out using a one-way analysis of variance (ANOVA) followed by a Dunnett multiple comparisons test. Data was scrutinized for normality using Kolmogorov-Smirnov (K-S) test. Statistical significance set a priori at p<0.05.

## Data Availability

Numeric data for this study can be found in the Supplementary Data File. The authors will make any other data, analytic methods, and study materials available to other researchers upon written request.

## Supplemental Information

**Supplemental Figure 1.**
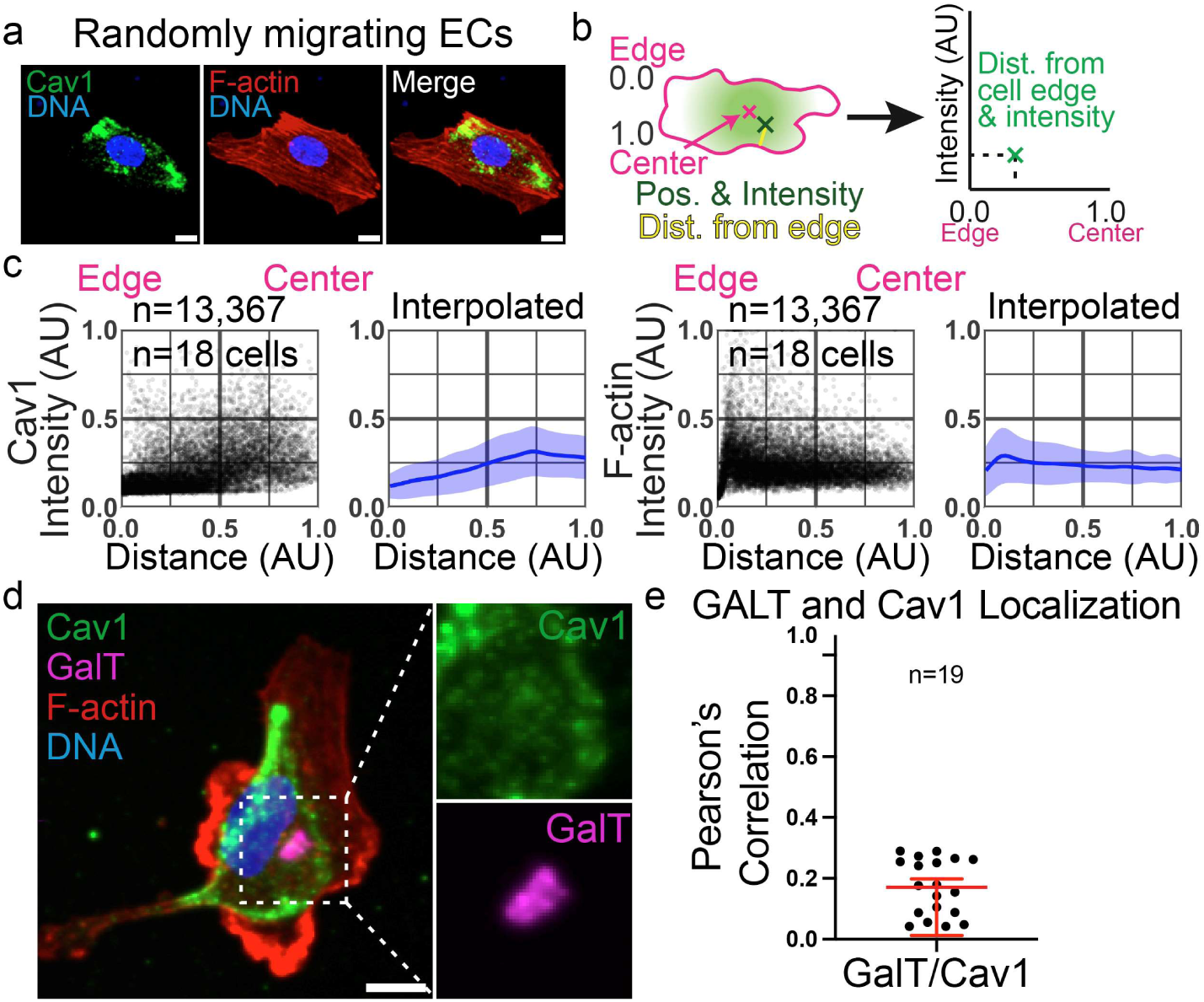
Caveolar organization in random cell migration. (**A**) Representative images of randomly migrating endothelial cells (ECs) fixed and stained for Caveolin-1 (Cav1, green), F-actin (red), and DNA (blue). Scale bars indicate 10 µm. (**B**) Cartoon describing the distance vs. intensity graphs shown in panel C. (**C**) Distance vs. intensity graphs showing quantification of distance from the cell edge (0) to the center of the cell (1) on the horizontal axis and normalized Cav1 or F-actin intensity on the vertical axis. Number of points and cells displayed on graph. (**D**) Representative images of an EC fixed and stained for Cav1 (green), F-actin (red), Galactosyltransferases (GalT, magenta), and DNA (blue). Scale bar represents 10 µm. Zoomed insets are shown to the right. (**E**) Graph displaying Pearson’s Correlation Coefficients from ECs comparing GalT and Cav1 colocalization. n = number of cells.

**Supplemental Figure 2.**
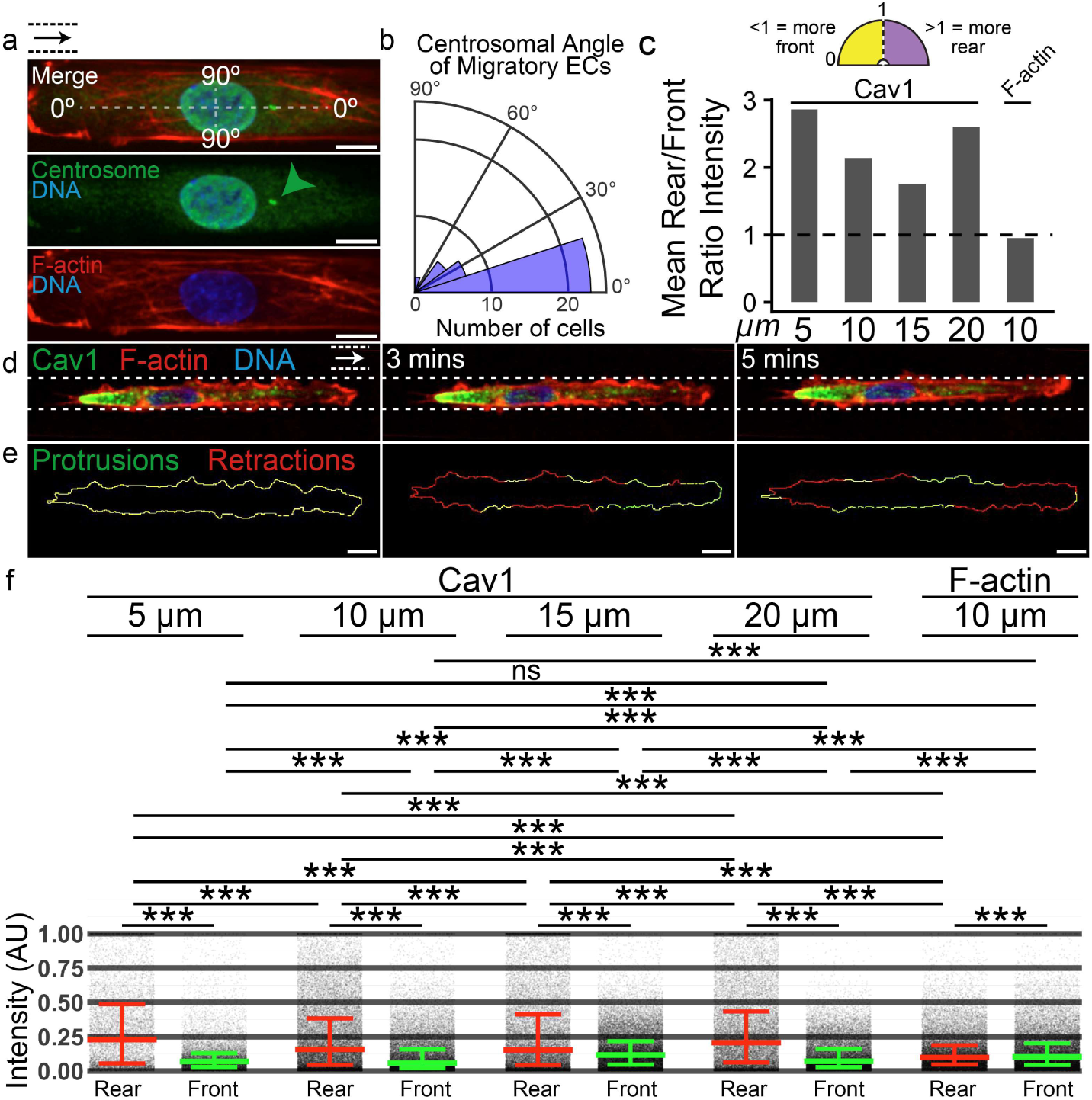
Line-patterned endothelial cells exhibit augmented front-rear polarity. (**A**) Representative images of individual endothelial cells (ECs) on line micropatterns fixed and stained for centrosome (green), F-actin (red), DNA (blue). Top image demonstrates the centrosomal position angles displayed in the graph of B. Green arrowhead in middle image indicates centrosome. Arrow in top left corner indicates the direction of cell migration. Scale bars indicate 10 µm. (**B**) Centrosomal angle graph corresponding to the images in A. (**C**) Mean Rear to Front ratio graph for Cav1 and F-actin. (**D**) Representative images from a movie displaying an endothelial cell on a line micropattern migrating from left to right at time courses 0 mins (left), 3 mins (middle) and 5 mins (right). (**E**) ADAPT plugin protrusion (green) and retraction (red) charts for the corresponding movie images above. Scale bars indicate 10 µm. (**F**) Cav1 and F-actin Rear vs. Front intensity plots. Box and whiskers indicate the median and interquartile range (IQR). *** indicates p-value from t-test of less than 0.001. ns = not significant (p-value > 0.05). Number of points per condition are as follows: 5 µm Cav1 Rear = 9,517; 5 µm Cav1 Front = 14,676; 10 µm Cav1 Rear = 28,589; 10 µm Cav1 Front = 33,031; 15 µm Cav1 Rear = 22,988; 15 µm Cav1 Front = 35,324; 20 µm Cav1 Rear = 21,026; 20 µm Cav1 Front = 29,872; F-actin Rear = 30,588; F-actin Front = 35,403. ***P<0.001. ns= non-significant.

**Supplemental Figure 3.**
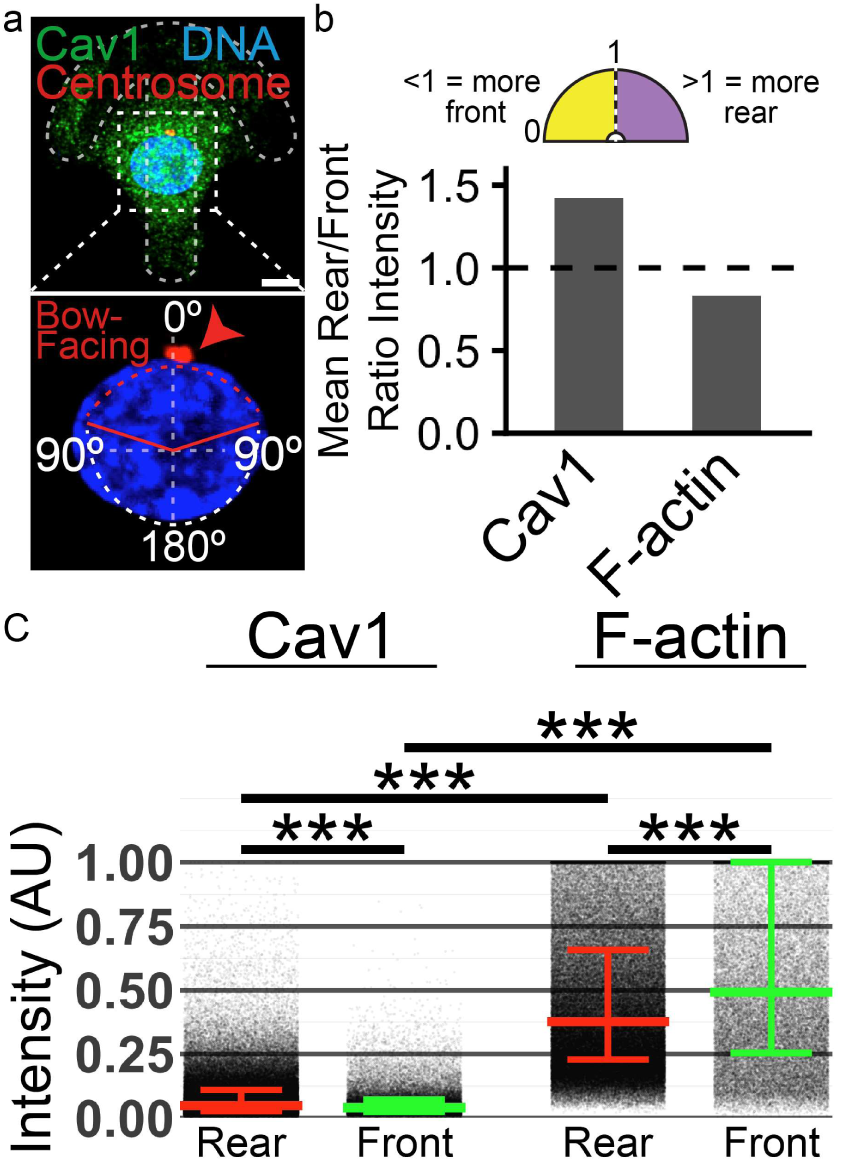
Crossbows planar polarized endothelial cells maintain caveolae in the rear. (**A**) Representative images of endothelial cells on crossbow micropatterns fixed and stained for Caveolin-1 (Cav1, green), Centrosome (red), and DNA (blue). Lower image, inset demonstrating the centrosomal position (indicated by a red arrowhead) relative to the crossbow’s orientation overlayed with a chart displaying corresponding angles. Scale bar indicates 10 µm. (**B**) Mean Rear to Front ratio graph for Cav1 and F-actin. (**C**) Cav1 and F-actin Rear vs. Front intensity plots. Box and whiskers indicate the median and interquartile range (IQR). *** indicates p-value from t-test of less than 0.001. Number of points per condition are as follows: Cav1 Rear = 142,755; Cav1 Front = 42,642; F-actin Rear = 146,036; F-actin Front = 43,878.

**Supplemental Figure 4.**
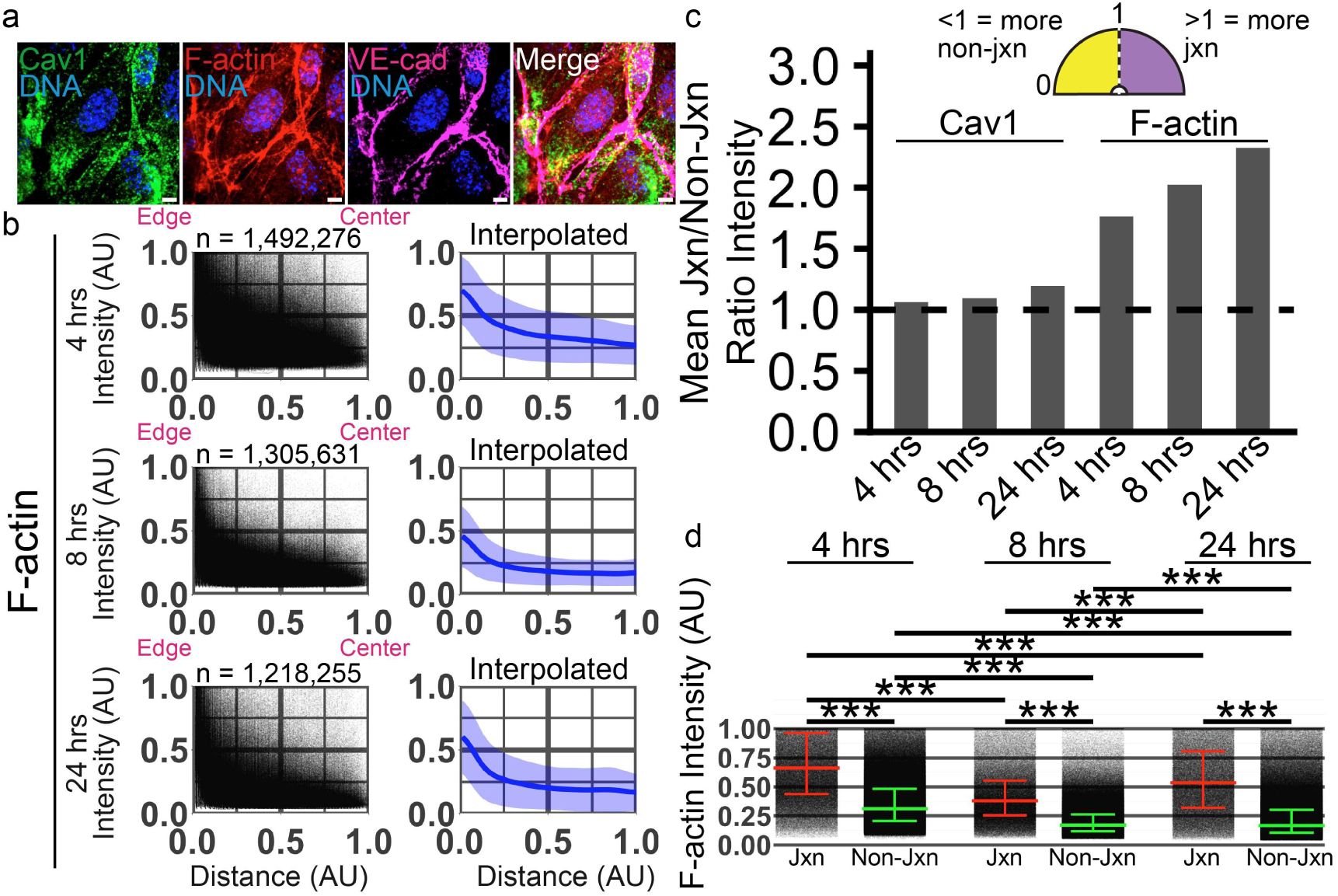
Caveolae are associated with junctions. (**A**) Endothelial cells fixed and stained for Caveolin-1 (Cav1 green), filamentous actin (F-actin, red), VE-Cad (magenta) and DNA (blue) in confluent imaging dishes. Scale bars represent 10 µm. (**B**) Distance vs. intensity graphs showing quantification of distance from the cell edge (0) to the center of the cell (1) on the horizontal axis and normalized F-actin intensity on the vertical axis. Number of points and cells displayed on graph. (**C**) Mean Junctional (Jxn) to non-junctional (Non-Jxn) ratio graph for Cav1 and F-actin. (**D**) F-actin Junctional vs. non-junctional intensity plots. Box and whiskers indicate the median and interquartile range (IQR). *** indicates p-value from t-test of less than 0.001. Number of points per condition are as follows: F-actin 4 hrs Junctional = 239,988; F-actin 4 hrs Non-Junctional = 1,252,288; F-actin 8 hrs Junctional = 208,030; F-actin 8 hrs Non-Junctional = 1,097,601; F-actin 24 hrs Junctional = 187,817; F-actin 24 hours Non-Junctional 1,030,438.

**Supplemental Figure 5.**
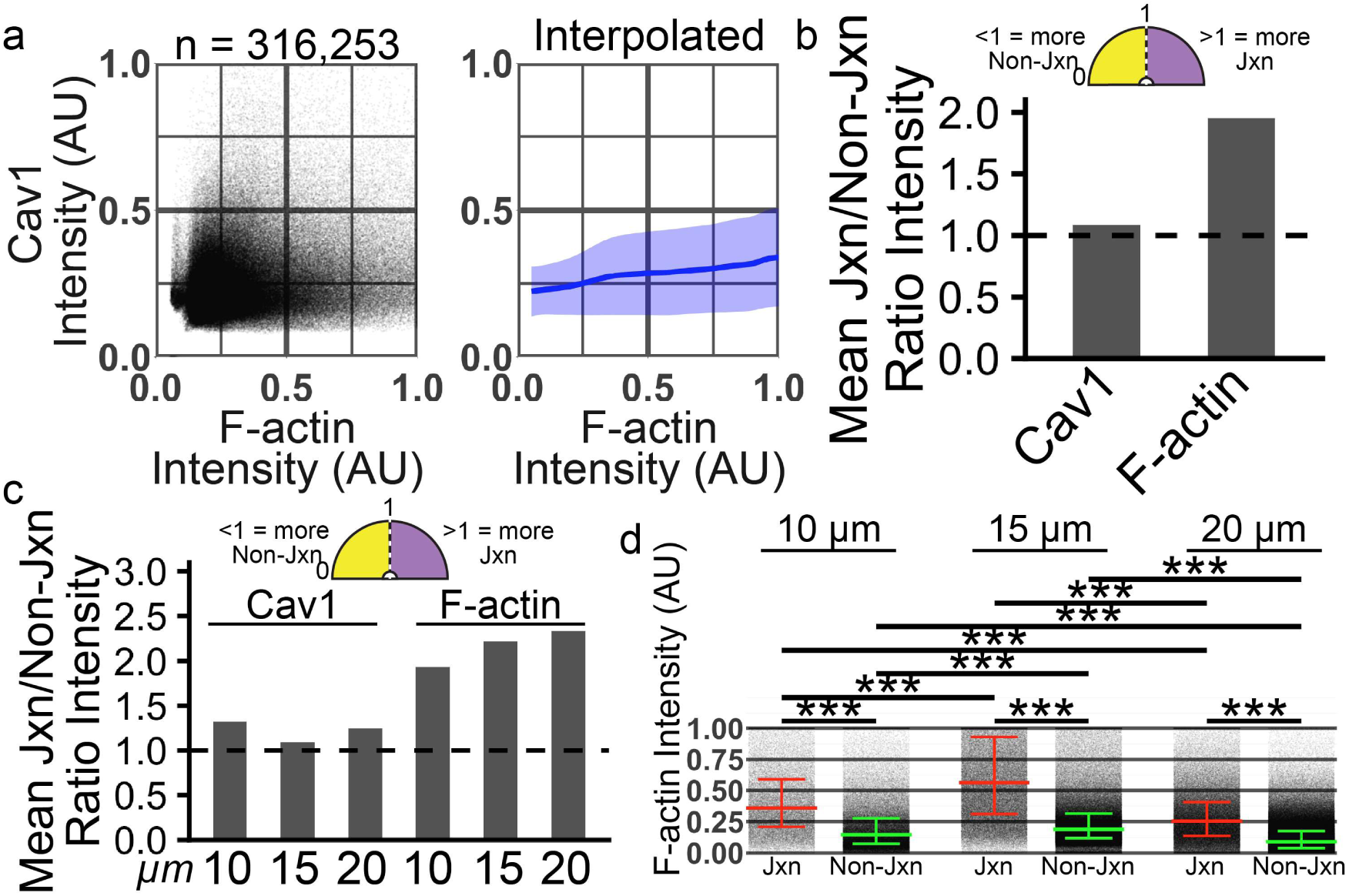
Influence of cell confinement on caveolae cellular localization. (**A**) Intensity plots and corresponding interpolation comparing Caveolin-1 (Cav1) and filamentous actin (F-actin). n=316,253. Intensity is measured using arbitrary units (AU). (**B**) Mean Junctional (Jxn) to non-junctional (Non-Jxn) ratio graph for Cav1 and F-actin for all lines only. (**C**) Mean Junctional to non-junctional ratio graph between differing line widths. (**D**) F-actin Junctional vs. non-junctional intensity plots. Box and whiskers indicate the median and interquartile range (IQR). *** indicates p-value from t-test of less than 0.001. Number of points per condition are as follows: F-actin Junctional 10 µm = 48,796; F-actin Junctional 15 µm = 120,416; F-actin Junctional 20 µm = 147,039; F-actin Non-Junctional 10 µm = 124,317; F-actin Non-Junctional 15 µm = 285,242; F-actin Non-Junctional 20 µm = 515,550.

**Supplemental Figure 6.**
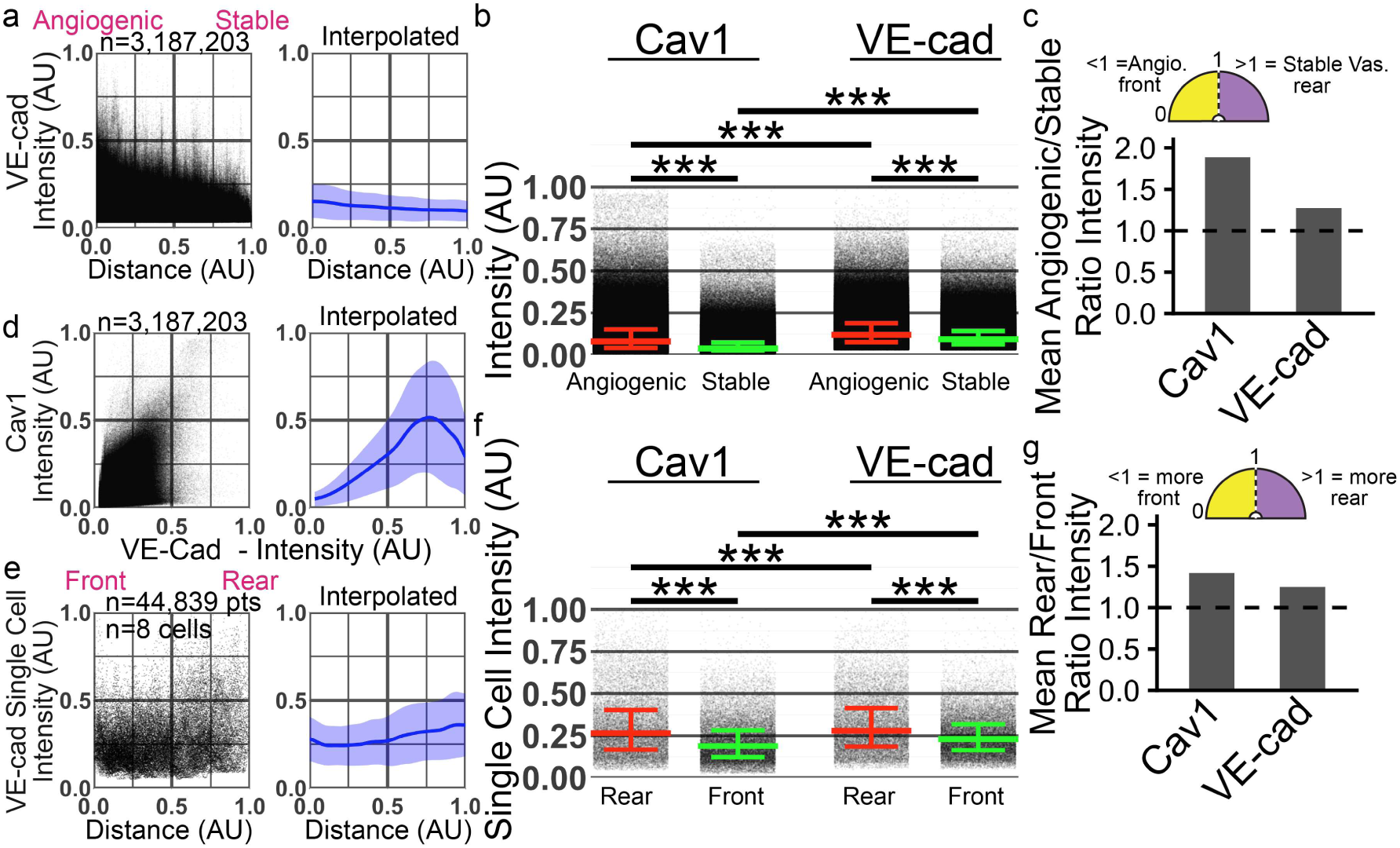
Caveolin-1 is upregulated at the angiogenic front in mouse retinal vasculature. (**A**) Distance from the angiogenic front (horizontal axis) vs. Normalized VE-cadherin (vascular endothelial cadherin, VE-cad) intensity (vertical axis). N = number of points. n = 3,187,203 pts collected from 10 images with an average of 54 cells per image. (**B**) Cav1 and VE-cad Angiogenic vs. Stable vasculature intensity plots, where angiogenic was within 25% of the maximal distance in the 60x image from the angiogenic front. Box and whiskers indicate the median and interquartile range (IQR). *** indicates p-value from t-test of less than 0.001. Points per condition are as follows: Cav1 Angiogenic Front = 1,457,069; Cav1 Stable = 1,730,134; VE-cad Angiogenic Front = 1,457,069; VE-cad Stable = 1,457,069. (**C**) Ratio of labeled protein intensities in the vascular plexus. (**D**) Intensity plots and corresponding interpolation comparing Cav1 and VE-cad from the angiogenic front images. n=3,187,203 pts collected from 10 images with an average of 54 cells per image. Intensity is measured using arbitrary units (AU). (**E**) Distance from the front of the cell (horizontal axis) vs. Normalized VE-cad intensity (vertical axis) plot. Number of points and cells displayed. (**F**) Cav1 and VE-cad Rear vs. Front single cell intensity plots, where rear was within 0.5 of the maximal distance in the 60x image from the angiogenic front. Box and whiskers indicate the median and interquartile range (IQR). *** indicates p-value from t-test of less than 0.001. Points per condition are as follows: Cav1 Rear = 17,915; Cav1 Front = 26,924; VE-cad Rear = 17,915; VE-cad Front = 26,924. (**G**) Ratio of labeled protein intensities in individual cells from the retinal vasculature.

**Supplemental Figure 7.**
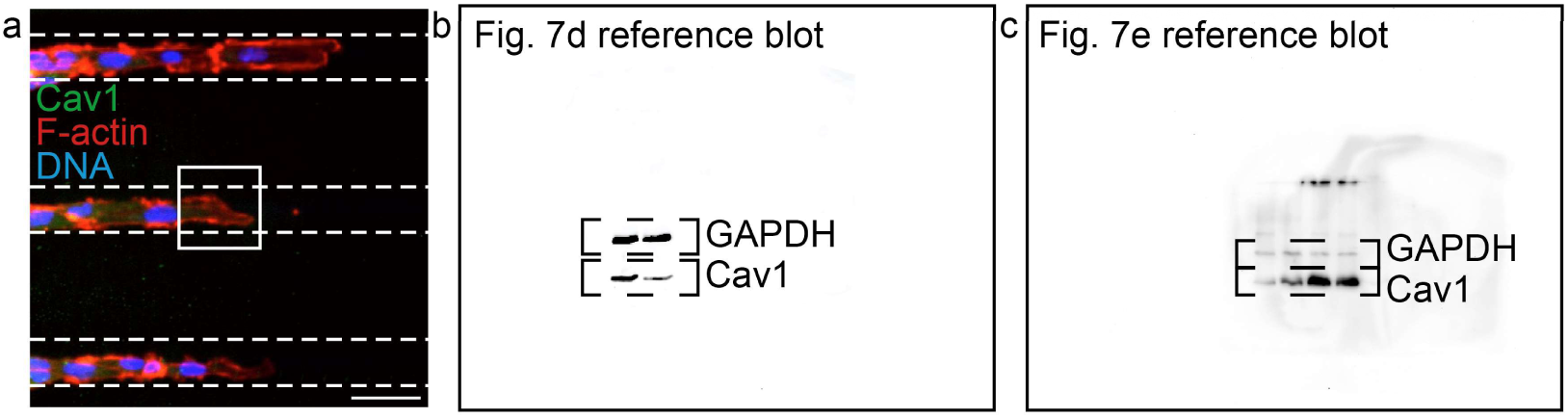
(**A**) Representative image of scratch wound line micropattern assay. (**B,C**) Western blots, uncropped.

**Supplementary Table 1.**
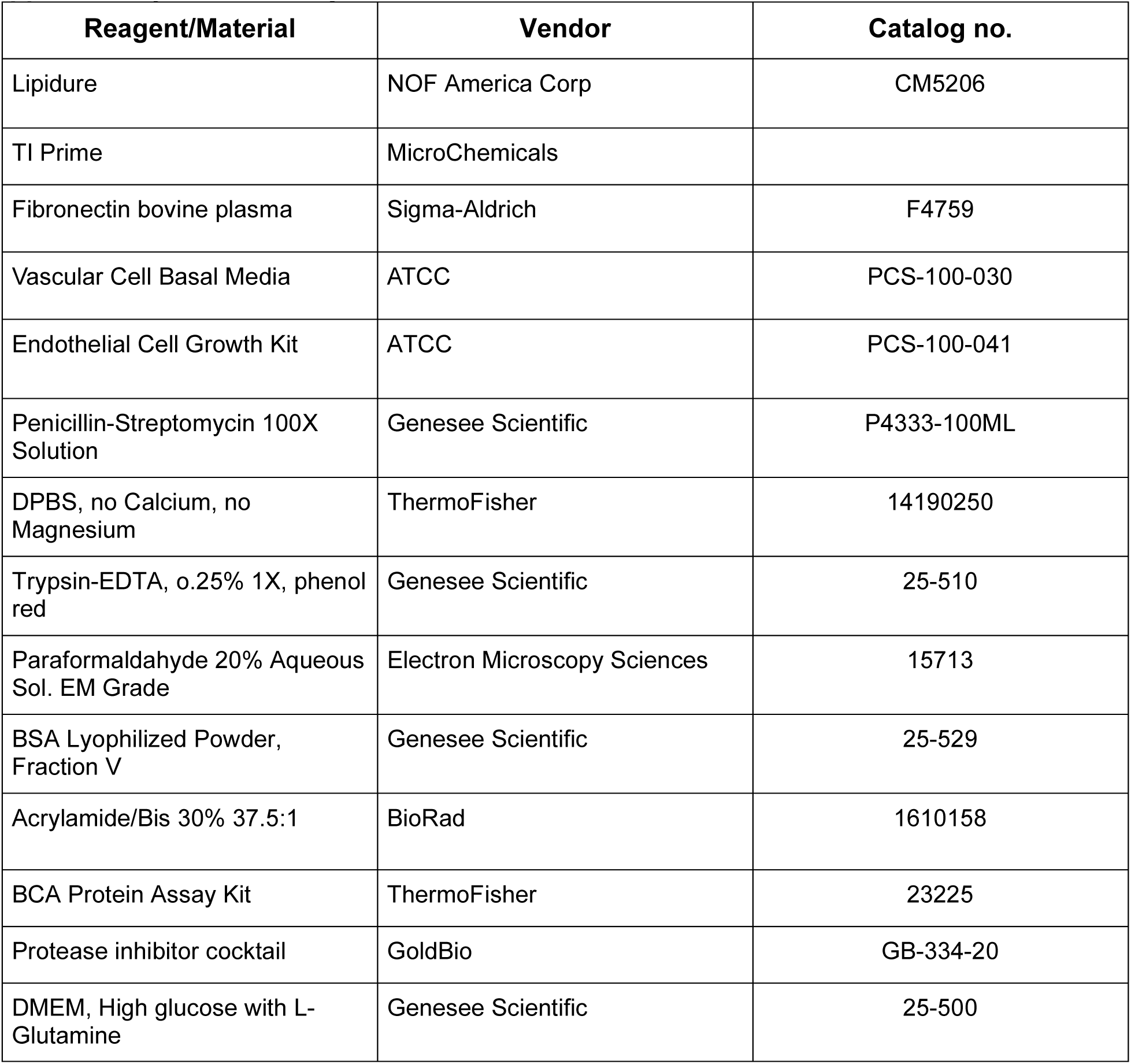
Reagents.

| **Reagent/Material** | **Vendor** | **Catalog no.** |
| --- | --- | --- |
| Lipidure | NOF America Corp | CM5206 |
| TI Prime | MicroChemicals |  |
| Fibronectin bovine plasma | Sigma-Aldrich | F4759 |
| Vascular Cell Basal Media | ATCC | PCS-100-030 |
| Endothelial Cell Growth Kit | ATCC | PCS-100-041 |
| Penicillin-Streptomycin 100X Solution | Genesee Scientific | P4333-100ML |
| DPBS, no Calcium, no Magnesium | ThermoFisher | 14190250 |
| Trypsin-EDTA, 0.25% 1X, phenol red | Genesee Scientific | 25-510 |
| Paraformaldehyde 20% Aqueous Sol. EM Grade | Electron Microscopy Sciences | 15713 |
| BSA Lyophilized Powder, Fraction V | Genesee Scientific | 25-529 |
| Acrylamide/Bis 30% 37.5:1 | BioRad | 1610158 |
| BCA Protein Assay Kit | ThermoFisher | 23225 |
| Protease inhibitor cocktail | GoldBio | GB-334-20 |
| DMEM, High glucose with L-Glutamine | Genesee Scientific | 25-500 |

**Supplementary Table 2.** Drug treatment compounds.

| **Name** | **Vendor** | **Catalog No.** | **Working Concentration** |
| --- | --- | --- | --- |
| VEGF | ThermoFisher | 100-20-10UG | 20ng/mL |

**Supplementary Table 3.** Antibodies.

| **Target Antigen** | **Vendor** | **Catalog No.** | **Working Concentration** |
| --- | --- | --- | --- |
| Cav1 Polyclonal | ThermoFisher | PA1064 | 1:1000 (WB) |
| Cav1 Monoclonal | ThermoFisher | 36000 | 1:1000(IHC) |
| GAPDH | ThermoFisher | PA1988 | 1:1000 (WB) |
| Centrin1 Polyclonal | ThermoFisher | 127941AP | 1:1000 (IHC) |
| VE-cadherin | Abcam | Ab33168 | 1:1000 (IHC) |
| Hoechst | ThermoFisher | 33342 | 1:1000 (IHC) |
| Alexa Fluor™ 647 Phalloidin | ThermoFisher | A22287 | 1µM (1:1000) |
| Alexa Fluor™ 555 Phalloidin | ThermoFisher | A34055 | 1µM (1:1000) |
| Goat anti-Rabbit IgG (H+L)<br>Secondary Antibody, Alexa<br>Fluor 488 | ThermoFisher | A11008 | 1µg/mL (1:1000) |
| Donkey anti-Rabbit IgG (H+L)<br>Secondary Antibody, Alexa<br>Fluor 555 | ThermoFisher | A31572 | 1µg/mL (1:1000) |
| Chicken anti-Rabbit IgG (H+L)<br>Cross-Adsorbed Secondary<br>Antibody, Alexa Fluor 647 | ThermoFisher | A21443 | 1µg/mL (1:1000) |
| Goat anti-Mouse IgG (H+L)<br>Highly Cross-Adsorbed<br>Secondary Antibody, Alexa<br>Fluor™ 488 | ThermoFisher | A11029 | 1µg/mL (1:1000) |
| Donkey anti-Mouse IgG (H+L)<br>Highly Cross-Adsorbed<br>Secondary Antibody, Alexa<br>Fluor™ 555 | ThermoFisher | A31570 | 1µg/mL (1:1000) |
| Goat Anti-Rabbit IgG (H+L) HRP | Genesee Scientific | 656120 | 1µg/mL (1:500) |

**Supplementary Table 4.** Plasmids.

| Name | Vendor | Catalog no. |
| --- | --- | --- |
| pmCherry-N1-GalT | Addgene | 87327 |

**Supplementary Table 5.** Micropattern materials.

| Device | Vendor |
| --- | --- |
| Photomask | Photomask portal |
| Spin coater model ws-640mz-23nppb | Laurell technologies |
| Plasma cleaner PE-25LF | Plasma Etch |
| JELIGHT deep uv model 24 | Jelight Company, inc. |
| Nikon eclipse TI Inverted microscope | Nikon |
| Laminar flow hood 1300 series A2 | ThermoFisher |
| Co<sup>2</sup> incubator MCO-170AICUVDL | Phcbi |
| Neon transfection system | ThermoFisher |

**Supplementary Table 6.** Statistics Values.

| Figure | Mean Normalized Intensity | Mean Difference<br>CI = Confidence Interval | Cohen's *d* coefficient<br>CI = Confidence Interval |
| --- | --- | --- | --- |
| **Supplemental Figure 2**<br>**(Single Cell Line Patterns)** | **<u>Caveolin-1:</u>** | **<u>Caveolin-1:</u>** | **<u>Caveolin1:</u>** |
|  | **<u>5 µm</u>** | **<u>5 µm Rear-Front:</u>** | **<u>5 µm Rear-Front:</u>** 0.93 |
|  | **Rear:** 0.3204486 | **Diff:** 0.2085667 | **95% CI:** (0.90, 0.96) |
|  | **Front:** 0.1118819 | **95% CI:** | **10 µm Rear-Front:** 0.64 |
|  | **<u>10 µm</u>** | **(0.2019353, 0.2151981)** | **95% CI:** (0.62, 0.66) |
|  | **Rear:** 0.251257 | **<u>10 µm Rear-Front:</u>** | **<u>15 µm Rear-Front:</u>** 0.54 |
|  | **Front:** 0.1171242 | **Diff:** 0.1341328 | **95% CI:** (0.53, 0.56) |
|  | **<u>15 µm</u>** | **95% CI:** | **20 µm Rear-Front:** 0.90 |
|  | **Rear:** 0.2719398 | **(0.1307017, 0.1375639)** | **95% CI:** (0.88, 0.92) |
|  | **Front:** 0.1546762 | **<u>15 µm Rear-Front:</u>** | **<u>F-actin:</u>** |
|  | **<u>20 µm</u>** | **Diff:** 0.1172637 | **Rear-Front:** -0.04 |
|  | **Front:** 0.2894802 | **95% CI:** | **95% CI:** (-0.06, -0.03) |
|  | **Rear:** 0.1115729 | **(0.1131920, 0.1213353)** |  |
|  | **<u>F-actin:</u>** | **<u>20 µm Rear-Front:</u>** |  |
|  |  | **Diff:** 0.1779074 |  |
|  | **Rear:** 0.1414351<br>**Front:** 0.1479031 | 95% CI:<br>(0.1739715, 0.1818432)<br><u>**F-actin:**</u><br>**Rear-Front:**<br>Diff: -0.006467977<br>95% CI: (-0.008665154, -0.004270799) |  |
| **Supplemental Figure 3 (Crossbow Patterns)** | <u>**Caveolin-1:**</u><br>**Rear:** 0.08260374<br>**Front:** 0.05805681 | <u>**Caveolin-1:**</u><br>**Rear-Front:**<br>Diff: 0.02464693<br>95% CI:<br>(0.02383767, 0.02525618) | <u>**Caveolin-1:**</u><br>**Rear-Front:** 0.3<br>95% CI: (0.28, 0.31) |
| **Figure 4 (Track Patterns)**<br>J = Junctional<br>NJ = Non-Junctional | <u>**Caveolin-1:**</u><br>**4 hours:**<br>J: 0.2032785<br>NJ: 0.1892599<br>**8 hours:**<br>J: 0.2114925<br>NJ: 0.1961792<br>**24 hours:**<br>J: 0.2991975<br>NJ: 0.2501426 | <u>**Caveolin-1:**</u><br>**4 hours J-NJ:** 0.01401868<br>95% CI:<br>(0.01327607, 0.01476130)<br>**8 hours J-NJ:** 0.01531335<br>95% CI:<br>(0.01413885, 0.01648786)<br>**24 hours J-NJ:**<br>0.04905488<br>95% CI:<br>(0.04753778, 0.05057198) | <u>**Caveolin-1:**</u><br>**4 hours J-NJ:** 0.12<br>95% CI: (0.12, 0.13)<br>**8 hours J-NJ:** 0.09<br>95% CI: (0.09, 0.10)<br>**24 hours J-NJ:** 0.25<br>95% CI: (0.25, 0.26)<br>**4-8 hours J-J:** -0.06<br>95% CI: (-0.06, -0.05)<br>**4-24 hours J-J:** -0.56<br>95% CI: (-0.57, -0.55)<br>**8-24 hours J-J:** -0.44<br>95% CI: (-0.45, -0.43) |
| **Supplemental Figure 4 (Track Patterns)**<br>J = Junctional<br>NJ = Non-Junctional | <u>**F-actin:**</u><br>**4 hours:**<br>J: 0.661821<br>NJ: 0.3747588<br>**8 hours:**<br>J: 0.4275565<br>NJ: 0.2112573<br>**24 hours:**<br>J: 0.5623166<br>NJ: 0.2419865 | <u>**F-actin:**</u><br>**4 hours J-NJ:**<br>0.2870622<br>95% CI:<br>(0.2859087, 0.2882157)<br>**8 hours J-NJ:**<br>0.2162992<br>95% CI:<br>(0.2152798, 0.2173185)<br>**24 hours J-NJ:**<br>0.3203301 | <u>**F-actin:**</u><br>**4 hours J-NJ:** 1.24<br>95% CI: (1.23, 1.24)<br>**8 hours J-NJ:** 1.40<br>95% CI: (1.40, 1.41)<br>**24 hours J-NJ:** 1.48<br>95% CI: (1.48, 1.49)<br>**4-8 hours J-J:** 0.93<br>95% CI: (0.92, 0.93) |
|  |  | 95% CI:<br>(0.3189784, 0.3216818) | **4-24 hours J-J:** 0.36<br>95% CI: (0.35, 0.36)<br>**8-24 hours J-J:** -0.52<br>95% CI: (-0.53, -0.52) |
| **Figure 5**<br>**(Confluent**<br>**Line Patterns)** | **<u>Caveolin-1 (5c):</u>**<br>J: 0.2809658<br>NJ: 0.2578761<br>**<u>Caveolin-1 (5e):</u>**<br>**10 µm:**<br>J: 0.4530674<br>NJ: 0.3423719<br>**15 µm:**<br>J: 0.2111021<br>NJ: 0.1941266<br>**20 µm:**<br>J: 0.1882475<br>NJ: 0.1509701 | **<u>Caveolin-1 (5c):</u>**<br>**J-NJ:**<br>Diff: 0.02308973<br>95% CI:<br>(0.02214451, 0.02403494)<br>**<u>Caveolin-1 (5e):</u>**<br>**10 µm J-NJ:** 0.1106955<br>95% CI:<br>(0.1077223, 0.1136688)<br>**15 µm J-NJ:** 0.01697545<br>95% CI:<br>(0.01599908, 0.01795181)<br>**20 µm J-NJ:** 0.03727741<br>95% CI:<br>(0.03661984, 0.03793498) | **<u>Caveolin-1 (5c):</u>**<br>**J-NJ:** 0.18<br>95% CI: (0.17, 0.19)<br>**<u>Caveolin-1 (5e):</u>**<br>**10 µm J-NJ:** 0.41<br>95% CI: (0.40, 0.42)<br>**15 µm J-NJ:** 0.12<br>95% CI: (0.11, 0.13)<br>**20 µm J-NJ:** 0.37<br>95% CI: (0.37, 0.38) |
| **Supplemental**<br>**Figure 5**<br>**(Confluent**<br>**Line Patterns)** | **<u>F-actin:</u>**<br>**10 µm:**<br>J: 0.4210111<br>NJ: 0.2183066<br>**15 µm:**<br>J: 0.5865579<br>NJ: 0.264581<br>**20 µm:**<br>J: 0.3020609<br>NJ: 0.129649 | **<u>F-actin:</u>**<br>**10 µm J-NJ:** 0.2027045<br>95% CI:<br>(0.2000122, 0.2053967)<br>**15 µm J-NJ:** 0.3219769<br>95% CI:<br>(0.3200199, 0.3239340)<br>**20 µm J-NJ:** 0.172412<br>95% CI:<br>(0.1712142, 0.1736097) | **<u>F-actin:</u>**<br>**10 µm J-NJ:** 0.87<br>95% CI: (0.86, 0.88)<br>**15 µm J-NJ:** 1.25<br>95% CI: (1.24, 1.26)<br>**20 µm J-NJ:** 1.09<br>95% CI: (1.09, 1.10) |
| **Supplemental**<br>**Figure 6**<br>**(Mouse**<br>**Retinal**<br>**vasculature)**<br>A = Angiogenic<br>S = Stable | **<u>Caveolin-1</u>**<br>**<u>Tissue:</u>**<br>A: 0.1143259<br>S: 0.6066319<br>**<u>Caveolin-1 Single</u>**<br>**<u>Cell:</u>**<br>Rear: 0.3087832<br>Front: 0.2178445 | **<u>Caveolin-1 Tissue:</u>**<br>**A-S:**<br>Diff: 0.05366273<br>95% CI:<br>(0.05347868, 0.05384579)<br>**<u>Caveolin-1 Single Cell:</u>**<br>**Rear-Front:**<br>Diff: 0.09093872 | **<u>Caveolin-1 Tissue:</u>**<br>**A-S:** 0.67<br>95% CI (0.67, 0.67)<br>**<u>Caveolin-1 Single Cell:</u>**<br>**Rear-Front:** 0.59<br>95% CI: (0.57, 0.61) |
|  |  | 95% CI:<br>(0.08783032, 0.09404713) |  |

